# Tissue and Extracellular Matrix Remodeling of the Subchondral Bone during Osteoarthritis of Knee Joints as revealed by Spatial Mass Spectrometry Imaging

**DOI:** 10.1101/2024.08.03.606482

**Authors:** Charles A. Schurman, Joanna Bons, Jonathon J. Woo, Cristal Yee, Nannan Tao, Tamara Alliston, Peggi Angel, Birgit Schilling

## Abstract

Osteoarthritis (OA) is a degenerative condition of the skeletal extracellular matrix (ECM) marked by the loss of articular cartilage and changes to subchondral bone homeostasis. Treatments for OA beyond full joint replacement are lacking primarily due to gaps in molecular knowledge of the biological drivers of disease. Mass Spectrometry Imaging (MSI) enables molecular spatial mapping of the proteomic landscape of tissues. Histologic sections of human tibial plateaus from knees of human OA patients and cadaveric controls were treated with collagenase III to target ECM proteins prior to MS Imaging of bone and cartilage proteins using a timsTOF fleX mass spectrometer. Spatial MSI data of the knee were processed and automatically segmented identifying distinct areas of knee joint damage. ECM peptide markers were compared between i) the medial halves of OA patient joints and the medial side of non-OA (cadaveric) joints, and ii) between the same medial OA tissues and their corresponding, less OA impacted, lateral joint halves. Distinct peptide signatures distinguished OA medial tissues from the cadaveric medial and OA lateral tissues (AUROC >0.85). Overall, 31 peptide candidates from ECM proteins, including Collagen alpha-1(I), Collagen alpha-1(III), and surprisingly, Collagen alpha-1(VI) and Collagen alpha-3(VI), exhibited significantly elevated abundance in diseased tissues. Highly specific hydroxyproline-containing collagen peptides, mainly from collagen type I, dominated OA subchondral bone directly under regions of lost cartilage. The identification of specific protein markers for subchondral bone remodeling in OA advances our molecular understanding of disease progression in OA and provides potential new biomarkers for OA detection and disease grading.

## Introduction

Osteoarthritis (OA) is the most common joint disease worldwide with the prevalence of OA increasing with age such that around half of all women (48.1%) and nearly one third of men (31.2%) aged 65 years and over ^1,2^ suffer from this debilitating joint disease. The future burden of musculoskeletal diseases is only expected to escalate in the coming decades as lifespans increase and more individuals become affected by comorbidities and other risk factors for OA, such as obesity, hypertension, dyslipidemia, dementia, and Alzheimer’s disease ^3,4^. Given that osteoarthritis and its risk factors all increase with age, we sought to understand more deeply the spatial progression of molecular changes throughout joint tissues in patients with clinically diagnosed OA and in non-arthritic individuals.

The relationship between subchondral bone and articular cartilage changes in OA, with well-documented progressive changes in subchondral bone remodeling, mineralization, vascularity, and innervation that parallel cartilage degeneration ^5,6^. In the early stages of OA, subchondral bone undergoes increased bone resorption, but at later stages, deregulated bone remodeling leads to increased subchondral bone density ^5–7^. In addition to the thickening of the subchondral bone plate, OA is often accompanied by microarchitectural changes in trabecular bone and bone marrow lesions ^8,9^. Indeed, clinical magnetic resonance imaging (MRI) identifies early changes to subchondral bone as predictors of later stage joint pain and joint replacement, even before detectable changes in the overlying cartilage ^10^. These observations motivate the search for diagnostics or interventions targeting subchondral bone to detect and retard OA progression, prior to the loss of articular cartilage, while the disease may still be reversible ^11^. However, many questions remain about the cellular and molecular targets for such interventions. Multiple cell types, including osteoblasts, osteoclasts, osteocytes, mesenchymal progenitors, and others, have been implicated in the subchondral bone changes in OA. In addition, aging, OA, and diseases such as type II diabetes cause changes in collagen post-translational modifications, including prolyl hydroxylation, glycation, and others^12,13^ tied to alterations in bone matrix material properties ^14,15^. In order to understand the loss of subchondral bone and cartilage homeostasis in OA, an unbiased spatial analysis of the proteomic changes in each component of the multi-tissue joint environment is needed. The complex nature of the ECM within bone and cartilage including their intrinsic properties have long obstructed proteomic profiling that has been successfully applied in most other tissues.

Matrix-Assisted Laser Desorption/Ionization - Mass Spectrometry Imaging (MALDI-MSI) is a rapidly advancing technology that allows for the direct detection of analytes in their spatial location directly from tissue sections. Initially introduced in the 1990s, MALDI-MSI has been applied in pathology research by providing an alternative to traditional histopathologic and immunocytochemical stains by directly imaging target proteins ^16,17^. Since its inception, MALDI-MSI has been used in several disease and tissue contexts including studies on neurodegenerative diseases and cancers, among others ^18–21^. This technology does not only comprise peptide-based spatial imaging but also MS Imaging of small molecule analytes, such as lipids, metabolites, and even large macromolecular glycan structures ^22–24^. The application of MALDI-MSI in the field of biomedical research, along with other mass spectrometry imaging technologies, such as water-assisted laser desorption ionization (SpiderMass)^25^ and parallel technologies like laser-captured microdissection^26^, opened new frontiers by enabling the identification of possible new diagnostic compounds or biomarkers in place in their direct biological microenvironment ^27^.

In previous years, investigators started to utilize MALDI-MSI to investigate the molecular landscape of OA, however, with a specific focus on lipids and glycans in cartilage and synovial tissues ^28–30^. Recently, MALDI-MSI was employed to investigate the lipid profiles and their distributions in the synovial membrane and infrapatellar fat pad of human OA patients, revealing the importance of lipid species in the development and regulation of inflammatory processes in OA, while others have extended these lipid and metabolite analyses to articular cartilage ^31–33^. Recently, spatial profiling of the subchondral bone has been achieved for lipids, expanding MALDI-MSI to mineralized tissues ^34–37^. Other investigators utilized MALDI-MSI to study N-glycans in the subchondral bone of the knee in OA and identified specific glycans within subchondral trabecular bone that were upregulated in different joint quadrants that they project correspond to areas of altered mechanical loading with disease ^30,38,39^. While the latter studies represent advancements in the use of spatial glycan imaging in subchondral bone, to date very few studies attempted imaging of <u>protein structures</u>^30,40,41^of joint tissues targeting the synovium and articular cartilage in human and equine samples.

The highly complex secondary and tertiary structure and the crosslinked nature of the bone ECM presents a technical hurdle in the application of MS Imaging for proteins and peptides to study subchondral bone changes in OA. However, recent significant advancements in the utilization of enzymatic protein digestions proved potent in the study of other ECM dense tissues and diseases such as breast, lung, and prostate cancers as well as in infarcted cardiac tissues ^20,21,42,43^. Collagenase III, also known as matrix metalloproteinase-13 (MMP-13), is a potent proteinase with a broad spectrum of activity targeting various ECM proteins, including collagen types I, II, and III. Homeostatically, this enzyme is involved in the degradation and turnover of the ECM, essential in processes such as skeletal development and remodeling, rendering it an ideal enzymatic choice to prepare bone and joint tissues for protein-based MALDI-MSI.

In this study, we spatially investigated the extracellular protein content and ECM remodeling of human end-stage OA of the knee, to demonstrate the first use of enzymatic digestion of formalin-fixed paraffin-embedded (FFPE) knee joint tissues (mounted on a microscopic ‘slide’) from non-arthritic cadaveric control donors (N=4) and age-matched OA patients (N=4) for MALDI-MS Imaging of ECM proteins (**Figure 1A**). In addition, these samples were prepared for serial glycan and ECM protein imaging ^44,45^, demonstrating the general ability to complete multi-omics imaging on the same slides of human joint tissues composed of both bone and cartilage regions. Our experimental strategy focused predominantly on spatial ECM imaging of bones and cartilage and allowed for the visualization and classification of differences in ECM protein composition in OA and during healthy aging, including differences in hydroxy-proline post-translational modifications. Through the identification of disease-regulated ECM proteins in OA, molecular mechanisms aggravating disease progression, potentially originating in the subchondral bone, may be spatially visualized for the first time to provide much needed novel disease markers that may be utilized to track, or even predict, osteoarthritis in humans.

**Figure 1:**
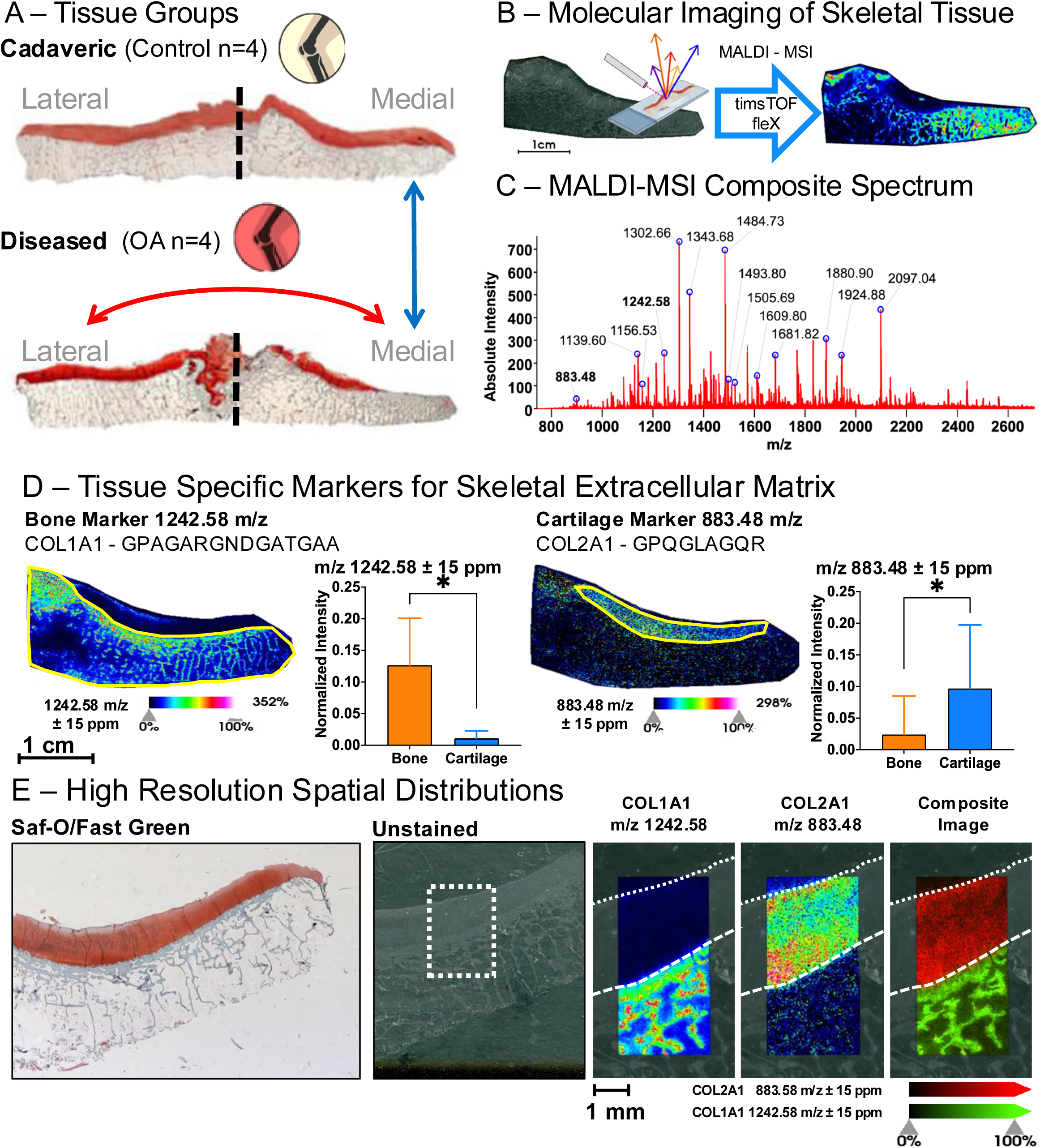
Arthritis of the Tibial Plateau and Spatial Proteomic Imaging. (A) Non-arthritic (control) cadaveric knee joint tissue donors without diagnosed arthritis, osteoporosis, or fracture, and Diseased Osteoarthritic (OA) patients receiving knee replacement composed a cohort that was age, sex, and BMI matched and stratified by their clinical OARSI (arthritis) grade. The Safranin-O/Fast Green histologic staining of the tibial plateau of a non-arthritic cadaveric donor (Control - top) and a representative patient with Osteoarthritis (OA - bottom) depicts the loss of proteoglycan staining (red) in Medial OA tissue, indicative of cartilage loss (lower right). (B) Molecular imaging of ECM proteins was accomplished with the timsTOF fleX (Bruker) to perform Matrix-Assisted Laser Desorption/Ionization (MALDI) Mass Spectrometry Imaging (MSI). (C) The composite MALDI-MSI spectrum was assembled from all imaged pixels from both medial and lateral halves of tibial plateaus from a single cadaveric (healthy) donor and an OA patient showed over 3,000 individual peaks. Subsequently, this allowed for the spatial extraction (spatial tissue mapping) of specific m/z features. (D) In a healthy, medial cadaveric tibial plateau with an intact cartilage layer, peptides of skeletal marker proteins Collagen alpha-1(I), for bone, (left) and Collagen alpha-1(II), for cartilage, (right) were preferentially detected in identified bone areas and cartilage areas (indicated by yellow outline) demonstrating the agreement between MALDI-MSI detected protein markers and the and known skeletal tissue region. Quantification is indicated with *p < 0.05 of average pixel intensity in bone (orange) vs cartilage (blue). (E) High-resolution scanning (20 micron laser step) in a tissue region composed of the bone and cartilage junction showed the clear spatial delineation of tissue type and spatial distribution of specific peptides within trabecular bone (m/z at 1242.58, COL1A1) and cartilage (m/z at 883.48, COL2A1).

## Results

### MS Imaging and Spatial Proteomics of Skeletal Tissues in the Knee

This study analyzed spatial molecular proteomic changes in the medial tibial plateau of the knee of a small prospective cohort of patients with medial osteoarthritis (OA) (N=4) compared to either the medial tibial plateau of non-arthritic cadaveric joints (control) (N=4) or the lateral tibial plateau of the same OA patients that did not show extensive cartilage loss via Safranin/Fast green histological stain (**Figure 1A**). This prospective OA cohort was composed of only male donors as were available through the Department of Veterans Affairs Medical Center (San Francisco, CA, USA). All donors received total knee arthroplasty at the Department of Veterans Affairs Medical Center. Non-arthritic control tissue was acquired through the Willed Body Program at the University of California San Francisco and selected to match OA donors in sex, BMI, and age resulting in a cohort of human tibial plateaus that were stratified by their Osteoarthritis Research Society International (OARSI) score (**Supplemental Figures 1** and **2, Supplemental Table 1**).

Given the complexity of osteoarthritis, and the need for improved molecular markers for the disease, we sought to utilize MALDI-MSI to investigate changes to the ECM in both subchondral bone and cartilage. Here, we demonstrate molecular imaging of proteomic data from knee joint tissues composed of both cartilage and bone regions using MALDI-MSI (**Figure 1B**). Given that MALDI-MS Imaging studies of collagen type proteins in bone have not previously been reported, we first verified that the enzymatic approach using collagenase III could successfully provide relevant details about the skeletal ECM. From a representative non-arthritic, cadaveric control, 33,234 individual MS1 mass spectra, one for each imaged spot, across the entire tissue section at a spatial resolution of 120 μm (laser step size). Averaging the spectra for all imaged pixels demonstrated a composite spectrum representative of the entire tissue (**Figure 1C**) with over 3,000 distinct individual peaks, each representing specific molecular features (peptides) with distinct mass-to-charge (m/z) ratios. Machine learning approaches on the top 500 most intense peaks were used to segment proteomes across the tissue (**Supplemental Figure 3**).

To determine how well the segmented proteins corresponded to actual tissue types, we investigated whether known protein markers of bone and cartilage would be preferentially located in these automatically identified regions (**Figure 1D**). Proteolytic peptides from Collagen alpha-1(I) and Collagen alpha-1(II), identified in studies of other tissue types ^20,42,46,47^, were used as markers for bone and cartilage tissue within the non-arthritic sample, respectively. The bone marker protein Collagen alpha-1(I) and its corresponding peptide G^317^PAGARGNDGATGAA^331^ with m/z at 1242.58, showed an intense signal predominantly in the ‘subchondral bone’ region (yellow outline, left), with almost no signal detected in the ‘cartilage’ region (yellow outline, right). Quantitatively, the pixel spots within the bone region had a significantly higher mean pixel intensity for the feature at m/z 1242.58 than the collection of pixels within the cartilage area by t-test (p<0.0001). Conversely, the cartilage marker protein Collagen alpha-1(II) and its peptide G^982^PQGLAGQR^990^ with m/z at 883.48, was most intensely abundant in the identified cartilage region with little to no signal intensity within the bone region. Compared via t-test, the generated cartilage region also showed a significantly elevated mean pixel intensity (p<0.0001) at this m/z compared to the bone regions. The segmented tissue regions corresponding to cartilage and bone also match well to histopathology via Safranin-O / Fast Green stains on serial sections of the tissues (**Supplemental Figure 2B**). To more clearly ascertain that MALDI-MSI can discern between bone and cartilage on the molecular level, a smaller region of the bone-cartilage osteochondral junction was imaged at an improved spatial resolution using a laser step size of 20 μm (**Figure 1E**), which (in our system) was the step size matching the minimal laser ablation area. At this improved higher spatial resolution, it was evident that the signal from the Collagen alpha-1(I) peptide G^317^PAGARGNDGATGAA^331^ at m/z 1242.58 emerged uniquely from the trabecular bone. In contrast the peptide from Collagen alpha-1(II) G^982^PQGLAGQR^990^ with m/z 883.48 was detected specifically in the upper cartilage region. **Figure 1E** highlights the distinct spatial distribution of the bone and cartilage markers, and the two areas display distinct separation when visualized in a composite image. To solidify this finding, additional unique peptides from Collagen alpha-1(I) and Collagen alpha-1(II), G^626^PAGERGEQ^634^ with m/z at 900.40 and G^999^PSGEPGKQGAP^1010^ with m/z at 1097.63 respectively, are also compared across the bone and cartilage regions and showed similarly significant differences in mean pixel intensity between bone and cartilage areas (**Supplemental Figure 4A-B**) providing distinct spatial features that correlate to the tissue types. These higher resolution settings (20 micron laser step size) revealed exciting spatial distributions for multiple features discernable with individual bone trabecula (**Supplemental Figure 4C**) – morphologic features that are themselves typically only at an average width of 200 microns. These spatial patterns reveal regions of high signal intensity specifically along the edges of trabecular bone for some m/z features, such as m/z 1943.91, while others show the most intense signal in the innermost portions of trabecula, e.g. at m/z 1242.58 and m/z 1406.71. While these specific peptide markers illustrate almost a binary delineation between bone and cartilage tissue regions, other unexplored markers such as glycans or other ECM proteins can reveal even further granularity or gradients across the bone-cartilage osteochondral junction.

Further imaging at higher resolution (20-micron laser step size), we sought to capture finer gradations of tissue compositional change across the osteochondral junction from the outer most cartilage surface deep into the trabecular bone. A well-known feature of the transition from cartilage to bone in the osteochondral junction is the calcified cartilage zone (CCZ)^48^ that lies in between the cartilage tidemark and the subchondral bone cement lines. These features were easily seen in histologic Safranin-O / Fast Green stains (**Figure 2A**). A thin strip of tissue, 2 mm in width, was scanned with MALDI-MSI from the cartilage surface to the bottom depth of trabecular bone in the tibial plateau at 20 μm laser step size. Segmentation in SCiLS within this region (**Figure 2B**) identified at least three distinct regions of cartilage including the outer articular surface layer (purple), internal cartilage (blue), and a lower outer boundary region separating cartilage from the subchondral bone (green) at the osteochondral junction, as indicated by the separation levels in the hierarchical clustering map. Subchondral bone separated into two distinct levels composed of an area of tissue containing the subchondral bone plate and adjacent trabecula (yellow) and a second region of deeper trabecular bone (orange). Discriminating feature analysis in SCiLS identified multiple m/z features that are characteristic of these different regions. For example, a marker specific to trabecular bone was found at m/z 1426.67 (**Figure 2C**) and a marker for cartilage was found at m/z 1480.75 (**Figure 2D**). Notably, features existed that had unique spatial distribution for the lower cartilage boundary region in between bone and cartilage at the osteochondral junction, such as the feature at m/z 1588.81 (**Figure 2E**). In order to fully appreciate the gradient of compositional change across the osteochondral junction, the spatial heatmaps for these selected m/z features are overlaid (**Figure 2F**), which shows a more gradual transition in molecular species through the tissue depth than the segmented regions alone. Investigating the osteochondral junction at higher magnitude (**Figure 2G**), the overlap of feature heatmaps revealed a region of tissue with strikingly similar delineation as the CCZ, where the most abundant intensity of the species at m/z 1588.81 appeared in an area matching the ‘cement line’ boundaries found in histological analysis. In addition, the species at m/z 1480.75 – a feature descriptive for cartilage – appeared in a region where the ‘cartilage tidemark’ was found histologically. In the future, further analysis and targeted LC-MS/MS will be needed to confirm the molecular identities of these additional specific molecular features. Currently, as we described above, the ability of our MALDI-MSI techniques to spatially visualize m/z features that uniquely match histologic features, even at small transitional areas like the calcified cartilage zone (CCZ), demonstrated the capability and premise of this technology for musculoskeletal biology.

**Figure 2:**
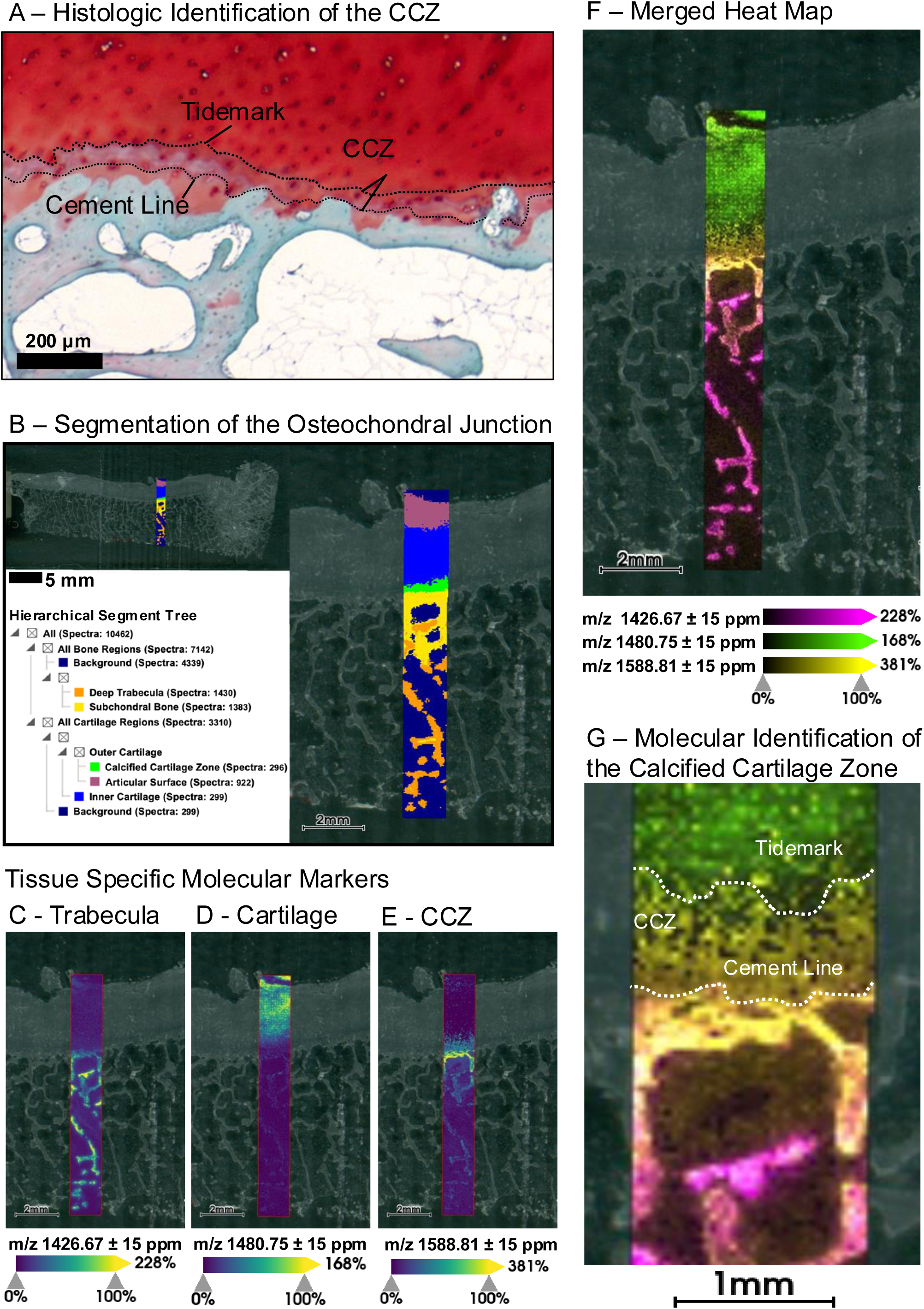
High Resolution MALDI-MSI of the Osteochondral Junction in Human Knees. **(A)** Safranin-O/Fast Green histologic staining of the tibial plateau at higher magnification reveals microstructural details at the osteochondral junction (OCJ) including the cartilage tidemark, cement lines, and the intervening calcified cartilage zone (CCZ). High resolution MALDI-MSI (20 micron laser step size) was completed from cartilage, across the OCJ, towards deep within the subchondral bone to visualize material differences across this tissue gradient. (B) Automated segmentation in SCiLS within this region identified several distinct tissue regions including multiple cartilage and subchondral bone layers, but also importantly multiple different regions around the OCJ (green, yellow). Discriminating Feature analysis found specific molecular features that described these new tissue regions including markers specific to trabecular bone (C) at m/z 1426.67 and for cartilage (D) at m/z 1480.75. (E) Specific features existed that revealed unique spatial distribution for the region in between bone and cartilage, such as the feature at m/z 1588.81. (F) Merging and overlaying of the spatial heatmaps for these three features revealed the gradient of changing matrix material composition through the OCJ. (G) Inspection of the OCJ directly using the combination of these molecular features revealed spatially distinct areas matching the location of cement line boundaries, the cartilage tidemark, and the CCZ as is seen in histologic staining.

### Subchondral Bone Remodeling in Osteoarthritis

Though some markers for cartilage degeneration can be detected in synovial fluid or in circulation, including small metabolites, lipids, inflammatory molecules, and by-products of proteoglycan and peptide catalysis^49^, less is known about markers for bone degeneration. Since we determined that MALDI-MSI can accurately identify tissue type and tissue specific protein markers in aged, non-arthritic bone and cartilage at high spatial resolution, we applied this technique to investigate the molecular landscape defining OA to potentially uncover novel markers of disease. We used a candidate-validation approach to first identify markers in a representative pairwise comparison between tissues for both the medial, non-arthritic cadaveric control group and the medial, OA cohort and then “validated” these markers using the full donor cohort. First, one medial, non-arthritic cadaveric tibial plateau was used as a control and was compared to one site-matched OA, or diseased, medial tibial plateau (**Figure 3A**). Acquired MALDI-MSI data from the representative control, C1, and the first OA patient, P1, were simultaneously loaded into the same data processing workspace. This resulted in a combined data set of 59,084 individual spectra. Automated region segmentation (**Figure 3B**) for the top 500 most intense peaks from the combined dataset resulted in identification of multiple tissue regions. Similar to the segmentation processes for control tissue, the segmented regions captured the cartilage regions and identified regions of non-arthritic subchondral bone in both control cadaveric tissue and in the deeper trabecular bone of the OA patient. Interestingly, this “non-arthritic” region existed in both groups, the cadaveric control and OA individuals. The non-arthritic regions in the OA patient indicated a molecularly more “healthy”-appearing bone region, that was distinct from other bone regions revealed in the right half, or outer portion, of the medial tissue from OA patient P1. In that specific area, where cartilage deterioration and arthritis damage were most severe, we identified a thin transitional layer of bone in the OA patient (**Figure 3B**, indicated in green) that separated the remaining healthier bone from a more profoundly impacted area of diseased arthritic tissue (**Figure 3B**, indicated in red).

**Figure 3:**
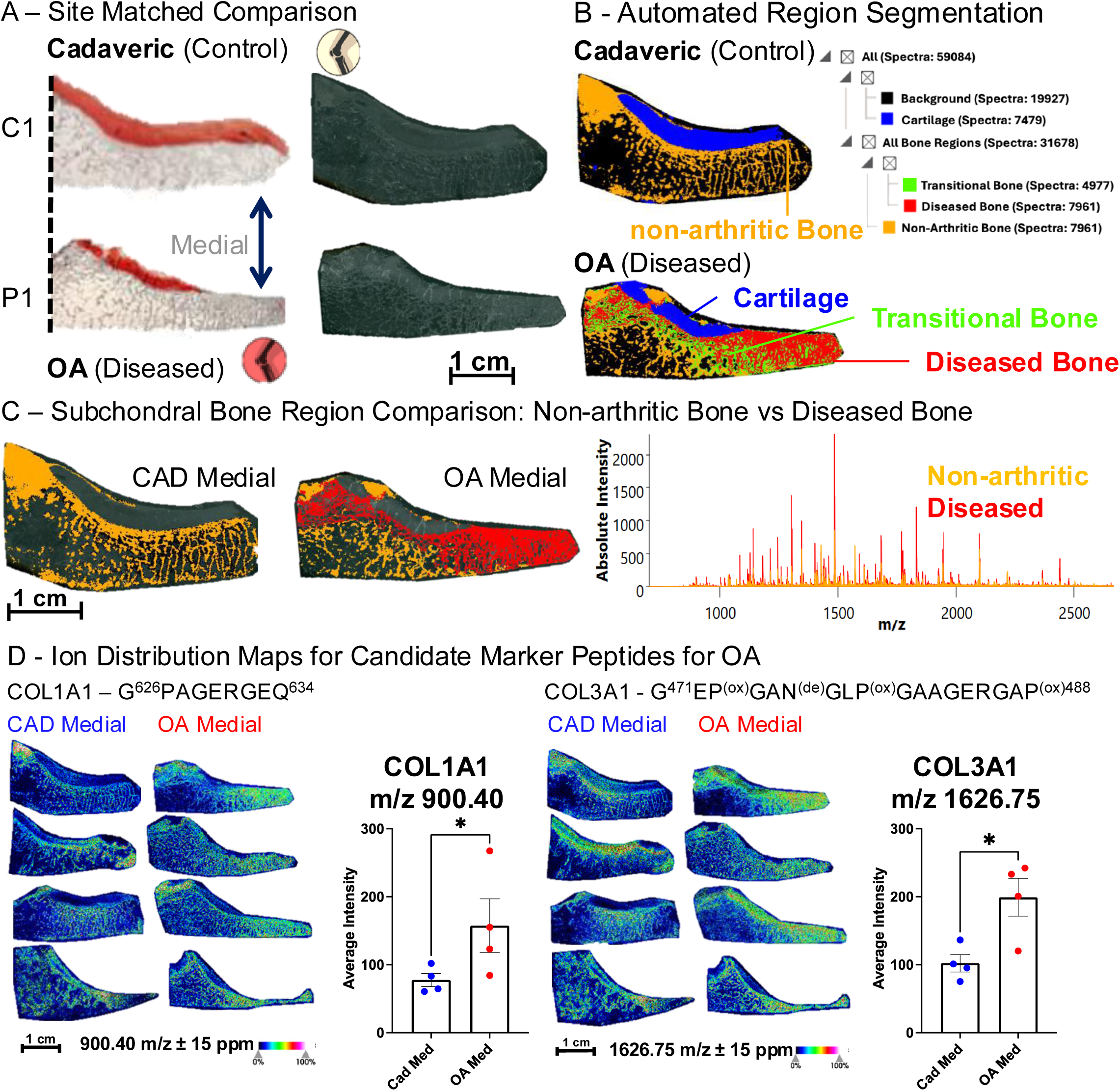
Site-matched Comparison, OA vs CAD Medial - Candidate Selection and Cohort Validation. (A) Tissue sections of medial Cadaveric (control) donors were compared against site-matched tissues from and medial OA (diseased) patients. (B) Automatic hierarchical segmentation was completed based on the top 500 m/z features from the combined datasets identifying cartilage (blue). Segmentation also identified 3 subtypes of bone including non-arthritic healthy bone (orange), a transitional layer (green), and a region of diseased bone (red) underneath the region of lost cartilage. (C) Comparison of the non-arthritic bone tissue (in orange) that were detected in both (cadaveric control and OA) with the diseased bone tissue (in red) that was detected in only the OA patient (right) showed that peak intensities for most molecular features in the MALDI-MS spectrum (right) were increased in the diseased tissue region. A pairwise ROC analysis was completed, and 31 unique molecular features were detected that illustrated the difference between the non-arthritic and diseased tissue types with a predictive AUROC of ≥0.85. (D) Several candidate features were isolated from the representative pairwise analysis and revealed significant differences between Cadaveric Medial (control) and OA Medial tissue sections within the full patient cohort using t-tests (n=4 per group), also see *Supplemental Table 2*. Examples of significantly increased disease markers by average pixel intensity from within the diseased tissue regions included a peptide from Collagen alpha-1(I) (left), G^626^PAGERGEQ^634^ at m/z 900.40, and a peptide from Collagen alpha-1(III) (right), G^471^EP^(ox)^GAN^(de)^GLP^(ox)^GAAGERGAP^(ox)488^ at m/z 1626.75. *p < 0.05, (n=4).

To investigate the molecular differences between all non-arthritic bone (**Figure 3B**, orange), including non-arthritic bone from the OA patient, and the OA diseased bone spatial regions (**Figure 3C**), and to generate candidate features for comparisons between the two groups, we applied a Discriminating Feature Analysis that utilized a receiver operating curve (ROC) scoring algorithm ^50,51^. The ROC curve analysis indicated how well the spatial distributions of individual m/z features matched or predicted the various bone regions and their spatial boundaries. We identified 31 candidate m/z features (**Supplemental Table 2**) that discriminated the non-arthritic and diseased spatial regions when requiring a statistical threshold with an ‘area under the receiver operating curve’ (AUROC) of >0.85. These m/z features represented the peptides that best differentiated between arthritic and non-diseased bone tissue across the representative tissue samples from cadaveric controls and OA patients. Candidate lists include peptides derived from collagen type I (Collagen alpha-1(I), Col1a1) and Collagen alpha-1(II), Col1A2) and collagen type III (Collagen alpha-1(III), Col3a1), see **Supplemental Table 2.**

To assess whether candidate features could serve as potential osteoarthritis markers, the mean pixel intensity of each feature (m/z) was used as a quantitative metric and compared from defined regions of tissue (**Supplemental Figure 5**) containing diseased subchondral bone from each individual OA patient (n=4) and non-diseased tissue from each cadaveric control (n=4). Signal from each candidate feature was exported for pixels within these ROIs to avoid the transitional bone and non-arthritic bone subtypes in OA samples and used to evaluate each candidate feature’s mean pixel intensity for each individual patient or control donor. The mean pixel intensity for each candidate feature was compared between the site-matched cadaveric controls and the medial OA samples to determine if there were robust differences across the patient cohort.

**Figure 3D** displays heatmaps for specific candidate m/z features that showed significant differences in abundance in OA (vs cadaveric non-arthritic) tissues. For example, the spatial distribution for the Collagen alpha-1(I) (COL1A1) peptide G^626^PAGERGEQ^634^ with m/z 900.40 (left panel) and the post-translationally modified Collagen alpha-1(III) (COL3A1) peptide G^471^EP^(ox)^GAN^(de)^GLP^(ox)^GAAGERGAP^(ox)488^ with m/z 1626.75 (right panel) illustrated statistically significant increases of each of these peptides in the medial portions of OA tissues in comparison to the medial cadaveric non-arthritic tissues. Overall, a signature of six candidate features, including peptides from Collagen alpha-1(I), Collagen alpha-1(II), FN1, and Collagen alpha-1(VI), was significantly regulated (p <0.05) comparing OA samples against cadaveric controls analyzing all four biological replicates per condition (N=4 OA, N=4 cadaveric non arthritic). Seven additional m/z candidate features, including peptides from Collagen alpha-1(III) and several novel features (without peptide/protein ID matches), trended towards significance (0.1< p <0.05) with OA **(Supplemental Table 2)**.

Joint degeneration occurs with age and can show differential severity on one side of the joint in OA. To model and visualize disease progression with age and OA, one candidate peptide from the ROC analysis, Collagen alpha-1(I) - G^305^LP^ox^GERGRP^ox^GAP^(ox)316^ with m/z 1211.61, was visualized and statistically quantified across both the lateral and medial halves of cadaveric ‘healthy’ donor C1 and OA patient P1 (**Supplemental Figure 6A**). While this Collagen alpha-1(I) feature was most highly elevated in the representative OA medial tissue (P1), the next highest intensity was observed in the medial side of the cadaveric control (C1) which was significantly higher than even in the lateral OA tissue (**Supplemental Figure 6B**). Therefore, MALDI-MSI detected age-related molecular OA signatures even within the cadaveric medial tissue group of ‘healthy’ controls. Even though age-related changes to OA-associated inflammatory markers and progression of articular cartilage damage markers are well known^52,53^, here we demonstrated underlying molecular and importantly spatial changes to the subchondral bone during joint disease progression with age. These findings warrant further investigation into age-related progression of OA in larger cohorts.

### Molecular Evidence of Localized and Site-Specific OA Bone Damage Preceding Cartilage Loss

Given the small sample size, the heterogeneity inherent to human patient samples, the many distinct endotypes of OA^54,55^, and natural age-related damage in the non-arthritic cadaveric joint cohort, analysis utilizing the lateral half of the OA joints as a patient matched control was justified to further investigate the molecular markers of OA found by MALDI-MSI (**Figure 4A**). Independent data sets for both halves of the tibial plateau from representative OA patient P1 were loaded into the same SCiLS data processing workspace to begin this new analysis following the same ‘candidate-validation’ approach used in the previous comparison. A composite spectrum was generated that represents the combined 64,183 individual spectra across the entire tibial plateau of OA patient P1 (**Figure 4B**). Region segmentation (**Figure 4C**) recapitulated similar bone regions found in the Medial OA section as before, including the non-arthritic medial subchondral bone (yellow) and a progressive change towards the most damage diseased bone region on the most medial edge (red). Interestingly, this diseased bone signature extended well into the lateral side of the joint where a thick cartilage layer remained and is not limited to the area of cartilage loss on the medial edge. The segmented regions containing the diseased bone signature and the healthier bone signature were separated out and a discriminating feature analysis utilizing the ROC prediction method was performed to generate candidate features with an AUROC > 0.85. A heat map for the example candidate peptide Collagen alpha-1(I) - G^626^PAGERGEQ^634^, with m/z at 900.40 and an AUC of 0.872, is shown beside its ROC curve to illustrate how well the distribution of the candidate features from this analysis aligns to the boundaries of the segmented regions (**Figure 4D**). Overall, 34 candidate features with an AUC > 0.85 (**Supplemental Table 2**), each with their spatial distributions, were found in this comparison to generate candidate molecular features for later cohort “validation” (**Figure 4E**). To verify that the candidate lists were not overly sample-specific, this same candidate feature identification process was completed on an additional OA patient (**Supplemental Figure 7**), and this resulted in a similar number of candidate marker features with the majority of features in both lists.

**Figure 4:**
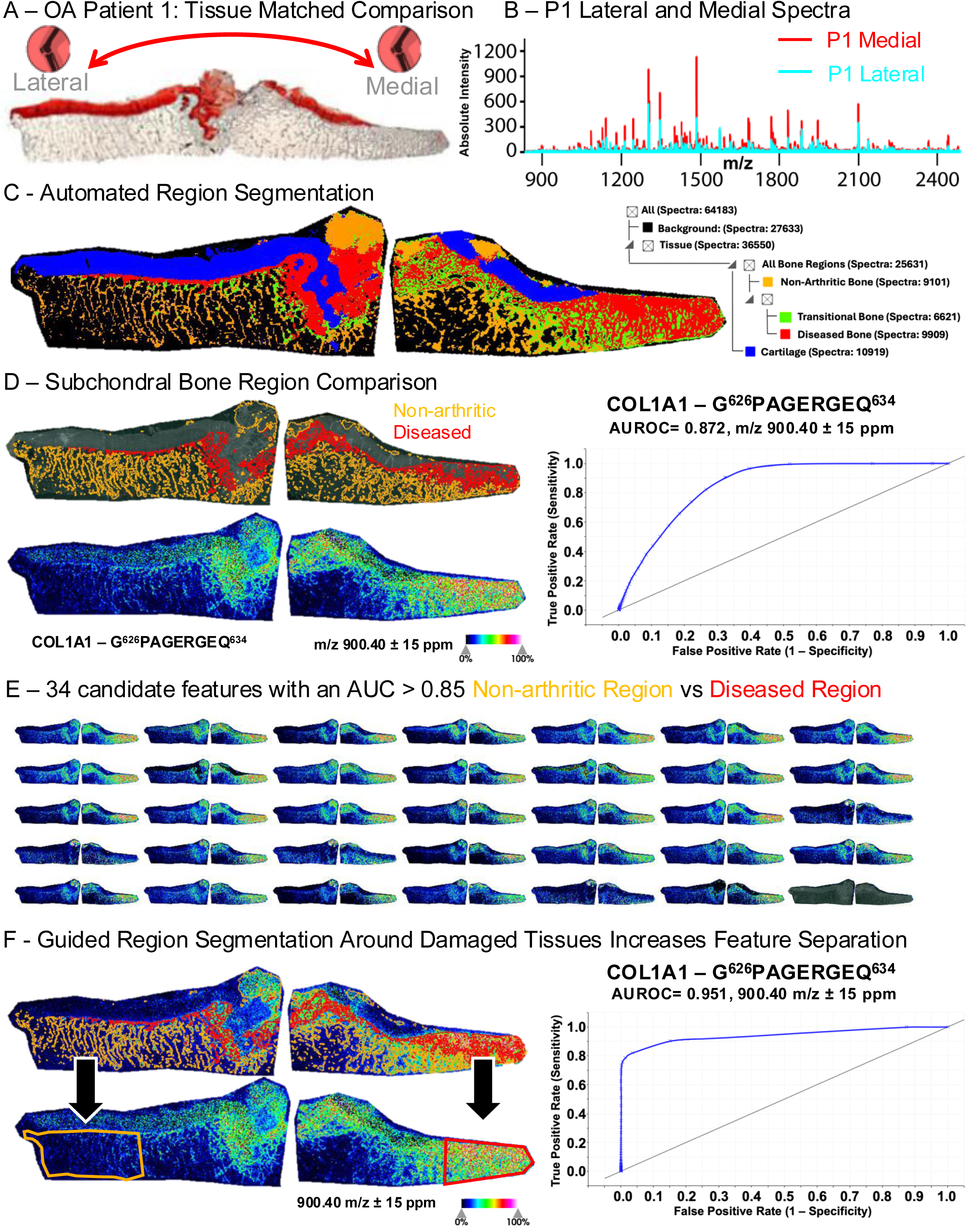
Tissue-matched Comparison, OA Medial vs OA Lateral. (A) Safranin-O staining of a representative arthritic tibial plateau from an OA patient (P1) showed the loss of cartilage (red) in the medial joint (right), however robust cartilage staining in the lateral joint was maintained (left). (B) MALDI-MSI data sets from the medial and lateral halves from the same patient P1 were combined into a single SCiLS Lab spectrum and peak intensities for molecular features were routinely higher in the OA medial section rather than in the OA lateral section. (C) Automatic hierarchical segmentation was completed based on the top 500 m/z features from both medial and lateral OA tissue sections identifying cartilage (blue) and bone regions including non-arthritic healthy trabecular bone (orange), a transitional layer (green), and a region of diseased bone (red) underneath the region of lost cartilage (showing sclerotic bone under areas of lost cartilage, red). (D) Pairwise comparison between the automatically generated subregions of the non-arthritic and diseased regions identified candidate features with AUROC ≥0.85. A representative peptide from Collagen alpha-1(I) (Col1a1), G^626^PAGERGEQ^634^, is shown next to its predictive ROC curve to illustrate how well candidate peptides’ spatial distributions were constrained to the identified “Diseased” region of the subchondral bone. (E) This analysis identified 34 different molecular features comparing Non-arthritic to Diseased bone regions in an individual OA patient. (F) Guided manual segmentation around the diseased regions of subchondral bone (up to the edge of exposed cartilage) on the medial side compared to the intact subchondral bone of the lateral joint increased the predictive power of the ROC analysis. The AUC for this peptide of Collagen alpha-1(I), G^626^PAGERGEQ^634^, at m/z = 900.40 increased from AUC = 0.872 to 0.951 with guided segmentation aided by both histology and the automated segmentation performed within SCiLS Lab.

Since the “diseased” region segment (red) that contains the diseased tissue appears somewhat underneath lateral cartilage and is not constrained solely to the medial tissue, we restricted comparisons to just the outermost regions of the medial and lateral joint to discriminate joint-site specific differences between diseased and healthier subchondral bone from the same donor (**Figure 4F**). New ROIs were generated using both histology and the automatically generated region segments as guides. Despite these new regions now containing multiple tissue types (marrow, trabecular bone) the predictive AUROC for many of the candidate features, including the depicted bone marker Collagen alpha-1(I) peptide, increased with the guided segmentation.

Finally, upon combination of both the candidate features from the initial comparisons of control cadaveric medial tissue to OA medial tissues (described in **Figure 3**) and the comparisons between OA lateral to and OA medial tissues (described in **Figure 4**), we obtained 44 unique candidate features or peptides that we subsequently assessed and ‘validated’ in the complete OA and cadaveric cohort (N=4 for each group). Guided segmentation was completed for the outer most edges of lateral and medial portions of each OA tibial plateau (**Figure 5A**). A PLS-DA based on the mean pixel intensity of all 44 candidate features from each tissue section (**Figure 5B**) showed that OA lateral samples cluster away from the OA medial samples despite them originating from the same donor. Next, individual candidate features were “validated” by the mean pixel intensity for the candidate across the 8 tissue sections (4 OA medial vs 4 OA lateral) with a paired t-test. Utilizing all 44 features for these comparisons gave a more comprehensive assessment of the molecular regulation occurring in OA. For instance, the candidate Collagen alpha-1(I) peptide G^305^LP^(ox)^GERGRP^(ox)^GAP^(ox)316^ with m/z 1211.60, initially only identified by comparing OA medial P1 to cadaveric medial C1, did not show significance on the cohort level in the earlier comparison of medial control cadaveric tissues to site-matched medial OA tissues (p = 0.229), but did show significant elevation in the OA cohort when comparing OA Medial to OA Lateral (p = 0.031) (**Supplemental Table 2**).

**Figure 5:**
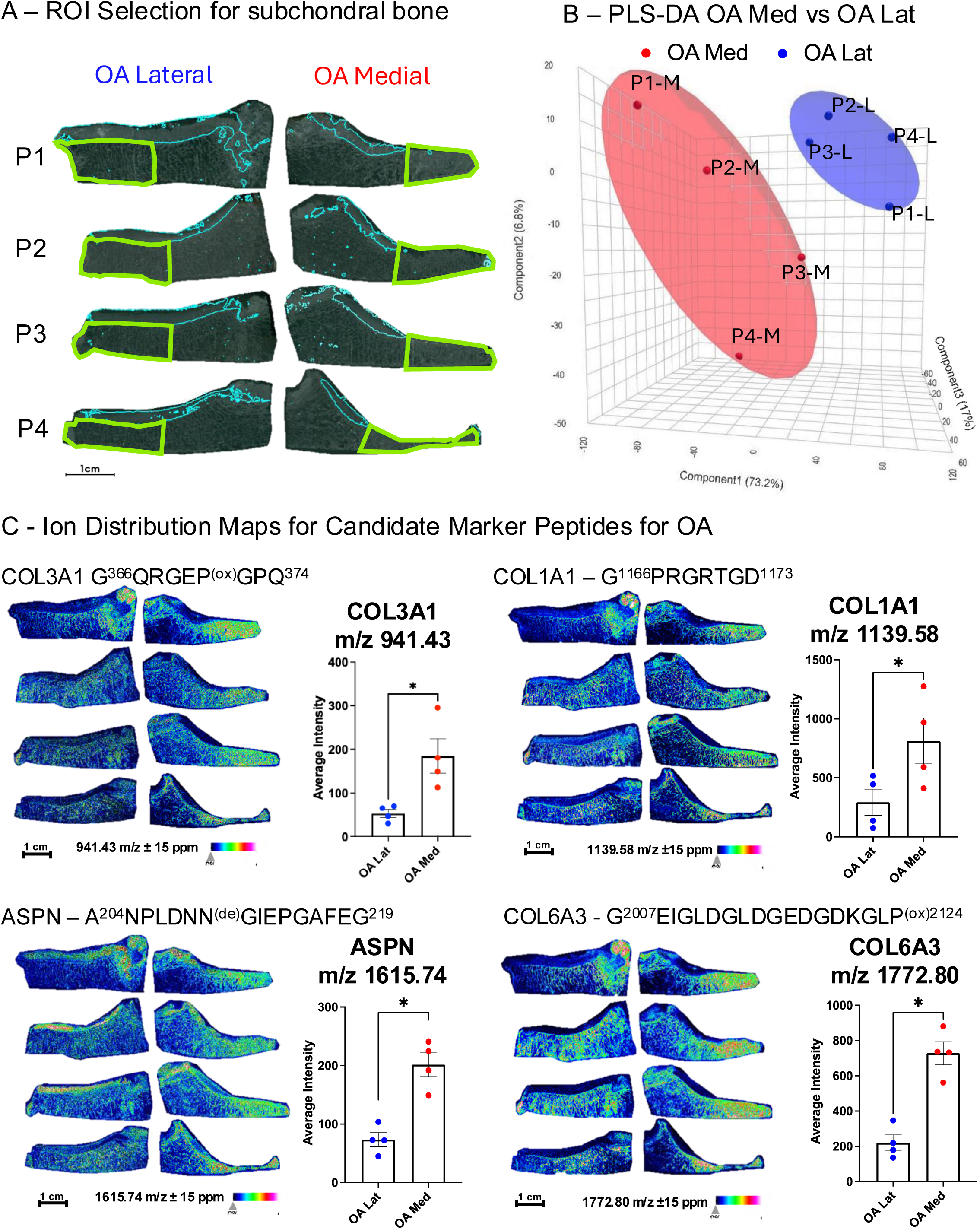
Candidate Ion validation Across Multiple OA Patients Comparing Medial to Lateral Joints. (A) Regions of Interest (ROIs in green) were manually segmented around the outer most regions of the medial and lateral tibial plateaus of OA patients (n=4), using the edge of the cartilage layer (thin blue line) as a guide for affected tissue in medial joints and using an equivalent distance on the lateral side. Average pixel intensity within the segmented region was calculated for each candidate ion that was previously identified using automated tissue segmentation and ROC curve predictive score. (B) Visualization of data grouping by 3-dimensional PLS-DA of pixel intensity showed distinct separation between the two joint sides despite physiologic complexity and patient heterogeneity in the OA medial cohort. (C) Group-wide heat maps of candidate feature distribution with peptide IDs derived from several ECM proteins or related proteins, Collagen alpha-1(III) (Col3A1) G^366^QRGEP^(ox)^GPQ^374^ at m/z 941.43, Collagen alpha-1(I) (Col1A1) G^1166^PRGRTGD^1173^ at 1139.58 m/z, Asporin (ASPN) A^204^NPLDNN^(de)^GIEPGAFEG^219^ at m/z 1615.74, and Collagen alpha-3(VI) (Col6A3) G^2007^EIGLDGLDGEDGDKGLP^(ox)2124^ at m/z 1772.80, depicted increases of the ion intensity in the outermost region of medial OA joints compared to their lateral counterparts (of the same OA patient). Statistical differences in feature intensities were confirmed via paired t-test *p < 0.05, (n=4) *see Supplemental Table 2*.

In statistical comparisons of the OA medial tissues vs their lateral tissue-matched controls using a paired t-test, 30 of the 44 candidate features showed significant differences, while an additional 8 features trended towards significance (**Supplemental Figure 8**). Of the 44 candidates, 36 m/z features matched to peptide identifications within 2 mDa of their theoretical mass from previous studies and our own spectral libraries generated ^20,42,46^ (**Supplemental Table 2**). Of the statistically elevated candidates in OA medial vs lateral tissues with known peptide identifications, four peptides belonging the fibrillar collagens Collagen alpha-1(III) and Collagen alpha-1(I), the network forming Collagen alpha-1(VI), and a component of the cartilage ECM, Asporin, that have been seen in proteomics of synovial fluid from OA patients (data not shown), are depicted (**Figure 5C**). Of the eight unidentified candidates, five reached significance levels in the group-wise comparison suggesting these may be novel bone-specific markers of OA that will require further classification, representing exciting possibilities for future worked concerned with subchondral bone degeneration in OA. Quantification of the full 44 candidates, with the additional 13 candidate features identified from this OA medial to OA lateral comparison, showed further differences in the earlier OA medial to control cadaveric medial comparison (**Supplemental Figure 9**). Overall, we demonstrate the capabilities of MALDI-MSI and an ECM-specific collagenase-based enzymatic digestion to detect osteoarthritis progression in the subchondral bone of human tibial plateaus and the potential to identify novel markers of disease.

### Spatial Distribution of Hydroxyproline Post-translational Modifications – Enrichment of Collagen Crosslinking in OA Subchondral Bone

One important mechanism in bone development and remodeling is the tightly regulated series of post-translational modifications that is required for the self-assembly of collagen fibrils (**Figure 6A**). This self-assembly requires the oxidation of specific proline residues along different collagen subtypes, which is catalyzed by several enzymes, including the family of prolyl hydroxylases ^56,57^. These homeostatic mechanisms are tightly regulated in healthy bone and joint tissues but have been implicated as dysregulated in different models of skeletal disease and aging in mice, including in diabetes and in disrupted osteocytic perilacunar/canalicular remodeling ^14,15^. Thus, the ability to detect specific post-translational modifications on distinct amino acids within a protein sequence from skeletal tissue, and the determination of their quantitative differences during disease or aging, will be a helpful tool to assess the molecular alterations occurring in aging and disease.

**Figure 6:**
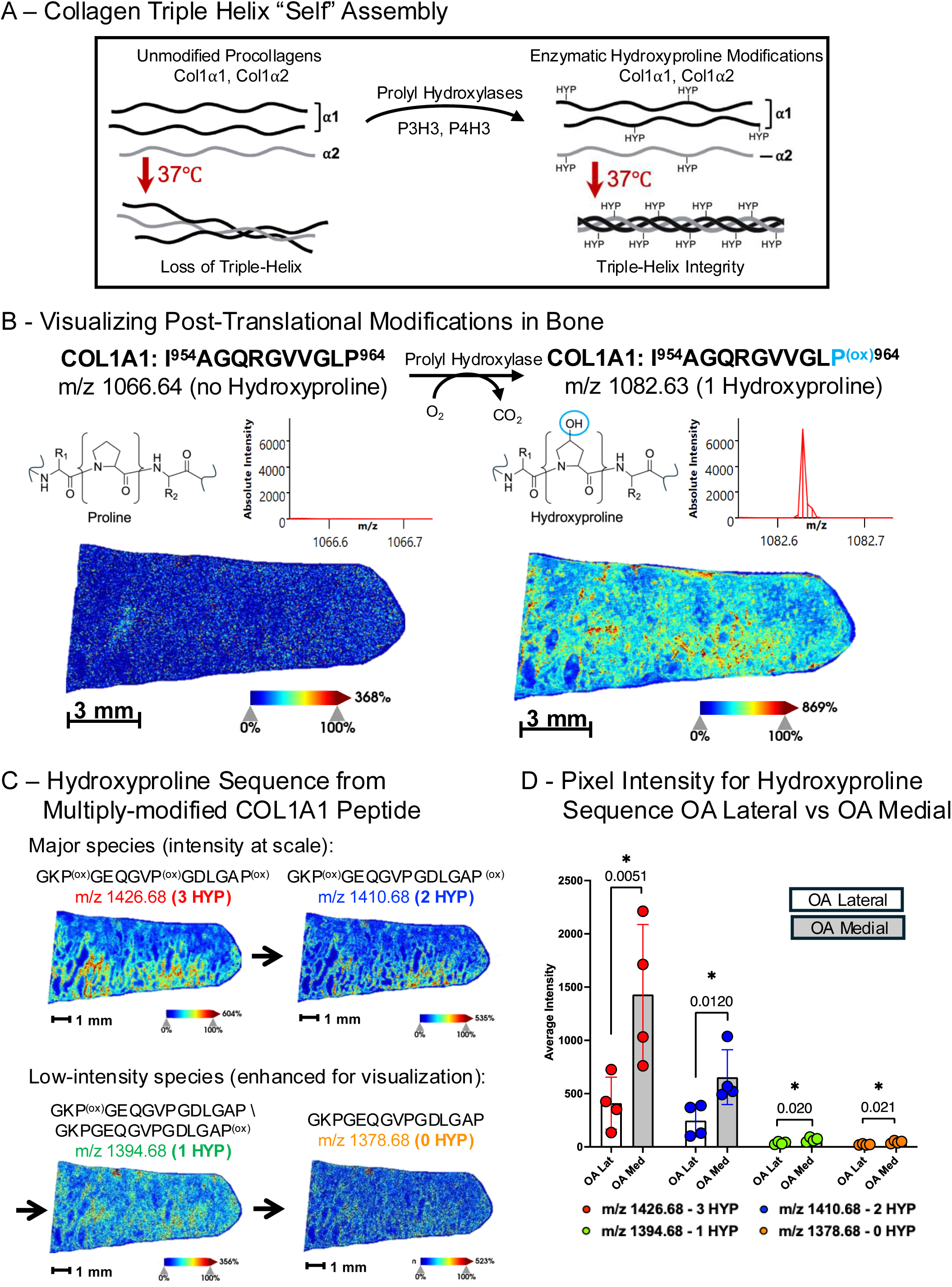
Post-translational Modifications of Collagens in OA Subchondral Bone. (A) Procollagen peptides are typically produced as unmodified amino acid chains. Without a series of post-translational modifications, including prolyl hydroxylation (via prolylhydroxylases), individual procollagens are unable to assemble properly into the tropocollagen triple helix, which is the basic protein building block of bone. (B) Heatmap of a previously identified unmodified peptide derived from Collagen alpha-1(I) (Col1A1), I^954^AGQRGVVGLP^964^, with m/z 1066.64 showed very low abundant (basically non-detectable) signal imaged at 20 micron laser step size in subchondral bone of osteoarthritic human tibial plateau (knee) FFPE tissue. Spatial imaging for the same peptide sequence with a modified hydroxyproline at Pro^964^, I^954^AGQRGVVGLP^(ox)964^ at m/z 1082.64, resulted in a well-defined spatial image with high intensity within subchondral bone trabecula. (C) The Heatmap of another Collagen alpha-1(I) peptide, G^656^KP^(ox)^GEQGVP^(ox)^GDLGAP^(ox)670^, identified at m/z 1426.68 with three hydroxyproline modifications is displayed (high intensity). The corresponding peptide sequence was also imaged for the peptide containing 2, 1, and 0 HYP residues, respectively, showing less abundant signal (at m/z 1410.68, 2 HYP) and lower quality spatial resolution (at m/z 1394.68 and m/z 1378.68, 1 and 0 HYP, respectively) as fewer HYP modifications were present in the target peptide. (D) Quantification of the signal intensity for this hydroxyproline-containing peptide series from subchondral bone comparing all replicates (N=4) from the Lateral (OA Lat, white bars) and Medial (OA Med, grey bars) halves for all OA patient tibial plateaus. * = p < 0.05 in paired t-test.

MALDI-MSI along with peptide spectral libraries from collagenase III digested tissues demonstrated the ability to detect and visualize matching peptides containing one or more oxidized proline residues in bone tissues. One peptide feature at m/z 1082.63 displayed very high signal intensity in the subchondral region below areas of cartilage loss in an OA patient (**Figure 6B**, right) and matched to a peptide from Collagen alpha-1(I) with a hydroxyproline (HYP) modification, I^954^AGQRGVVGLP^(ox)964^, with a small mass error (<15 ppm). We also visualized the spatial distribution of the corresponding peptide without HYP modification (unmodified proline) I^954^AGQRGVVGLP^964^ with a precursor ion at m/z 1066.64 (**Figure 6B**, left), and the latter unmodified peptide was not detected above noise level indicating no discernable bone trabecular morphology, while the HYP modified peptide was highly abundant. To confirm the identity of the modified HYP peptide we performed liquid chromatography tandem mass spectrometry (LC-MS/MS) data-dependent acquisitions (DDA) using the timsTOF HT (Bruker) of collagenase digested tissues recovered from slides after imaging confirmed the identity of the modified peptide and the specific site-location of the modified proline, Pro^964^ in the peptide IAGQRGVVGLP^(ox)^ of Collagen alpha-1(I) on the MS/MS level (**Supplemental Figure 10A**). A precursor for the unmodified peptide, I^954^AGQRGVVGLP^964^, was not detected in this LC-MS/MS experiment consistent with the MALDI MSI data (described above). The presence of the modified peptide I^954^AGQRGVVGLP^(ox)964^ and the absence of the corresponding unmodified form emphasized a first case of a dramatic shift in proline modifications during OA and the potential of these peptides to discriminate mature or diseased skeletal tissues.

This integrated workflow of combining MALDI-MS Imaging and ESI-MS/MS spectral library building was used to determine which modified forms of peptides - specifically hydroxy proline-containing peptides - were more abundant with disease or OA. For example, another feature at m/z 1426.68 displayed an abundant spatial profile with localized areas of high intensity near the center of the bone trabecula in a region of sclerotic OA subchondral bone (**Figure 6C**). This feature (m/z 1426.68) matched to a triply-modified peptide of Collagen alpha-1(I), G^656^KP^(ox)^GEQGVP^(ox)^GDLGAP^(ox)670^. A feature at m/z 1410.68, corresponding to the same peptide sequence with one less HYP, G^656^KP^(ox)^GEQGVPGDLGAP^(ox)670,^ presented with a similar spatial distribution as the triply-modified peptide including similar regions of signal intensity within the central areas of trabecular bone. LC-MS/MS DDA confirmed the presence of peptide precursors in our data set with three (**Supplemental Figure 10B**) and two HYP residues (**Supplemental Figure 10C**) for this peptide. MS/MS spectra for the doubly HYP modified peptide were able to distinctly localize the position of the two HYP modifications to prolines Pro^658^ and Pro^670^.

Attempts to assay this peptides sequence with yet fewer post-translational modifications, such as the same base peptide with 1 HYP (at m/z 1394.68), and the unmodified peptide (at m/z 1378.68), resulted in low abundant signals in the MALDI MSI spectra (high signal to noise). Additionally, precursor ions for these species were not detected in the corresponding LC-MS/MS acquisitions and generated spectral library. Comparisons investigating the OA cohort (**Figure 6D**) for this Collagen alpha-1(I) peptide in all its HYP-modified forms, starting with the triply modified form G^656^KP^(ox)^GEQGVP^(ox)^GDLGAP^(ox)670^, with m/z at 1426.68 and the doubly modified form at m/z 1410.68 (2x HYP) were highly abundant and <u>significantly upregulated in the</u> <u>diseased OA tissue</u> (OA medial-grey bar vs OA lateral – white bar). We also extracted pixel intensity for corresponding peptides with only 1 HYP at m/z 1394.68 or 0 HYP at m/z 1378.68, however their average pixel intensities were extremely low, not reaching above the surrounding signal-to-noise (S/N) background, and they were observed at almost 30 times lower intensity than the triply-modified peptide. Overall, we determined that collagen-derived peptides containing (multiple) HYP modifications were more prevalent in bone material and - more importantly - can become over-represented in OA and disease tissues vs control (healthy) tissues.

## Discussion

Osteoarthritis is most commonly first indicated by self-reported pain in patients, however, also by the appearance of osteophytes, or mineralized growths, and by joint-space narrowing as revealed by X-rays or Magnetic Resonance Imaging (MRI) scans of affected joints. Even then, full confirmation or grading of the disease state cannot be made until histopathologic stains that can confirm the loss of cartilage and other markers of disease are visualized on post-surgically removed joint tissue. Further, there are currently only limited pre-clinical molecular diagnostics for OA or OA progression that stratify patients who may benefit from early intervention to improve joint health. In this study we demonstrate that matrix-assisted laser/desorption ionization (MALDI) mass spectrometry imaging (MSI) may be used as an effective tool for the study of the molecular and biochemical spatial changes that occur in human joint tissues during osteoarthritis (OA) and that molecular insight gained from these studies can inform development of non-invasive diagnostic biomarkers for disease progression.

We report the first successful protein-based molecular spatial mapping of skeletal tissues composed of both cartilage and bone regions by targeting ECM proteins with an enzymatic strategy utilizing collagenase-III. The feature-rich composite spectrum of collagenase III digested skeletal tissue captured by MALDI-MSI molecularly discriminates bone from cartilage, demonstrating the effectiveness and promise of this technology to study musculoskeletal disease and its effects on the complexity of the extracellular matrix (ECM). Proteins detected comprise standard ‘marker’ proteins for bone and cartilage, including Collagen alpha-1(I) and Collagen alpha-1(II) respectively, spatially localized to their corresponding tissue types as confirmed by histology. Additionally, higher resolution imaging at the border between bone and cartilage identifies specific molecular features with spatial distributions that can recapitulate features found through traditional histopathologic approaches. Molecular image processing allowed for parallel assaying of over several hundred candidate peptide features at once, enabling a comprehensive workflow for unbiased detection of novel disease markers. In FFPE tissue sections from the tibial plateau of total knee replacement patients and site-matched tissues from non-arthritic, age, and BMI matched cadaveric donors, we find specific peptides belonging to ECM proteins that are upregulated in OA. These include peptides belonging to less common collagens such as Collagen alpha-1(III) and Collagen alpha-3(VI). The identified panel of peptide markers was expanded by utilizing the less affected lateral half of joints from the same OA donors as patient and tissue matched controls. This limited person-to-person variability and allowed for the validation of over 30 specific peptides that were significantly upregulated in medial OA tissues compared to either cadaveric medial or OA lateral controls. Interestingly, in our analysis, no candidate features were significantly down regulated in OA subchondral bone among our small clinical cohort. This could be an effect of the small cohort size or the targeted use of collagenase III to investigate matrix protein changes, or our primary focus on bone markers in end-stage disease conditions. Still, the identification of protein level regulation in the subchondral bone of OA suggests that molecular remodeling on the protein level occurs throughout the process of joint degeneration in addition to alterations to bone density and bone mass. Knowledge of how different peptide regions of ECM proteins are altered in disease provides needed biologic context for the mechanical and biochemical changes occurring in OA diseased tissues that may directly influence resident cell types, as well as the potential ability to use these peptides as therapeutic or diagnostic biomarkers in future studies. This study provides the successful application of MALDI-MSI for protein-based imaging on composite human joint tissues including both cartilage and bone, as well as biologic insight into joint disease on the protein and molecular level which can inform future studies aimed at improving treatment and diagnostics for OA.

A significant insight offered by this OA imaging study is the direct visualization of the spatial distribution of ‘diseased’ subchondral bone on the protein level. While it is intuitive that damage and associated changes to the ECM would be most highly constrained to areas of subchondral bone underneath lost cartilage, proteomic profiling of altered bone ECM composition has not before been achieved. These changes primarily occur in areas where osteocyte dysfunction and subchondral bone sclerosis have been described histologically and radiographically (**Supplemental Figure 1**), as demonstrated in previous studies on this cohort of patients ^7^. While this disease signature is most evident in areas underlying cartilage loss, imaging of the lateral half of OA joints reveals that the disease signature emerges even in subchondral bone areas where a thick cartilage layer remains (**Figure 4C red, Figure 5C**). While osteocyte dysfunction plays a causal role in OA, alterations to subchondral bone also result from perturbations to MSC, osteoblast and osteoclast function; thus it will be important to further investigate the cellular origins of this altered proteomic signature^7,58^. Such studies may elucidate the extent to which progression of OA arises from altered joint mechanical loading, changes in cellular metabolic or secretory behavior, or altered signaling by resident skeletal or immune cell populations. Since the loss of subchondral bone homeostasis can precede cartilage damage, the search for early OA diagnostics should focus on bone. Thus, the continued investigation into subchondral bone changes, especially at earlier ages, among a larger cohort of individuals is warranted. Additionally, given that the current OA cohort was limited to male patients enrolled in the Veteran’s Administration healthcare system, and that OA has an increased prevalence in females^1,2^, proteomic profiling of OA in female joints is essential and should be prioritized in future studies.

Of the many exciting findings from the molecular imaging of subchondral bone in osteoarthritis, one major outcome was the identification of multiple collagen species within the disease signature of subchondral bone. Here, an enzymatic pretreatment of the tissues was completed with collagenase III to target the ECM and ECM-associated proteins. Due to the nature of collagenase III, it is not surprising that many members of the candidate features identified in this study are a form of collagen. Collagenase III, or MMP13, is already known to be active and contribute to OA related damage^7,59^, and so additional application of this enzyme to the tissue may have masked potential endogenous or existing damage marker peptides. However, given that the dramatic increase in signal for damage marker peptides is localized to sclerotic regions of subchondral bone under cartilage loss and not in areas of remaining subchondral bone exposed to the same enzymatic pretreatment process, this enzymatic processing accurately allowed for the identification of broad molecular changes occurring in subchondral bone with OA.

Here we provide molecular evidence of bone remodeling in OA on the protein level through the regulation of several collagen subtypes in bone. The regulation of these specific collagens and ECM proteins in disease can additionally give insight into biologic mechanisms of disease. Collagen itself is not simply a structural protein, for example it is well known that the collagen fibril is bioactive and can bind many other proteins or provide Arginine-Glycine-Aspartic (RGD) domains for cellular attachments^60–62^. In a specific example, the unmodified Collagen alpha-1(I) peptide G^1165^PRGRTGD^1173^ detected at m/z 1139.58 is significantly increased in the diseased medial regions of our OA patients (**Figure 5C**). This specific peptide exists towards the C-terminal domain of the collagen 1 triple helix and lies within an area of interleukin 2 (IL2) and amyloid precursor protein (APP) binding ^61,62^ (**Supplemental Figure 11A**). IL2 is a well-known inflammatory marker and enhancer of the T-cell immune response, while APP interaction with collagen 1 has also been shown to enhance cell-matrix adhesion and enhance the function and activation of peripheral monocytes ^63,64^. Thus, the detection of this specific peptide in the subchondral bone of OA can point towards molecular mechanisms by which inflammation may be amplified in OA. This peptide also overlaps an area of Phosphophoryn, also known as Dentin phosphoprotein - a protein known to enhance mineralization in the formation of dentin tissue ^65^. The implication of proteins involved in mineral regulation in OA-affected tissue could provide mechanistic insight into the deregulation of mineralization and subchondral bone thickness that occur in OA, though these interactions remain to be investigated. These novel observations in the context of OA and knowledge of protein spatial distribution specifically in the subchondral bone beneath areas of cartilage loss provide new insights into the cellular and molecular interactions that may be occurring in disease and promote further investigation of the biologically regulatory features of ‘structural’ collagen in the context of this disease.

Other collagen I peptides identified as upregulated in OA when compared against non-arthritic, cadaveric controls or histologically normal lateral joint tissues, are those identified with post-translational modifications. Hydroxyproline modifications in bone are important enzymatically generated modifications that increase collagen structural integrity and reinforce the triple helix of the collagen fibril ^57,66^. The upregulation of these modifications in OA suggests molecular mechanisms impacting the material stiffness of bone in OA joints independently of hypermineralization. Further examples of PTM regulation in OA can be found in other upregulated regions of the collagen type 1 fibril. For example, the peptide of Collagen alpha-1(I) G^626^PAGERGEQ^634^ at m/z 900.40 within the mature collagen type I fibril is bound to a region of Collagen alpha-1(II) that contains lysine Lys^343^ - a known glycation site (**Supplemental Figure 11B**) ^62^. This region of the mature collagen fibril is also thought to bind different keratin sulfate proteoglycans (**Supplemental Figure 11B**), both important PTMs for collagen ^61,62^. Another peptide directly identified with HYP modifications and upregulated in OA, G^565^KPGEQGVPGDLGAP^670^ also contains a documented glycation site at Lys^657.^ Glycation, in the form of advanced glycation end-products (AGEs), can further stiffen the bone matrix ^67^ and alterations to proteoglycans in bone, such as in diseases like osteogenesis imperfecta, negatively impact bone strength ^68^. 22 of the 27 significantly elevated candidate features identified as belonging to Collagen alpha-1(I) or Collagen alpha-1(II) contained glycation or other known locations of collagen post-translational modifications that are not ascribed to HYPs when compared to the functional collagen interactome (**Supplemental Table 2**). Thus, the presence of these modification sites in OA bone where cartilage has worn away implicates post-translational regulation as one mechanism that may alter subchondral bone homeostasis in OA. Interestingly, we find that these PTM changes occur in areas of cartilage loss where it is thought that physical loading of the joint alters throughout the course of disease ^69^. It remains to be elucidated whether these changes reflect the mechanical reinforcement of collagen by PTMs in bone underlying areas of cartilage damage or are a mark of underlying disease in bone that gives rise to local cartilage degeneration.

Aside from collagen 1, the presence of other less common collagens comprises the detected disease signature of OA related peptides in bone. Several peptides of Collagen alpha-1(III), a less common fibrillar collagen active in wound repair, is upregulated in the diseased subchondral bone of OA patients (**Figure 5C**). Collagen Type III has also been implicated in the cartilaginous regions of osteoarthritic joints as an attempt by chondrocytes to remodel or stabilize remaining cartilage ^70^, while in bone it is thought to promote osteoblast differentiation leading to increases in trabecular bone mass ^71^. Thus, the upregulation of Collagen Type III in the subchondral bone of OA tissue compared to control tissue points to a potential role in driving late stage OA increases to bone mass and trabecular bone sclerosis common in OA ^72^.

Another less common collagen type detected in OA subchondral bone through MALDI-MSI was Collagen Type VI. Though commonly seen in muscle and tendon, in the pericellular matrix around chondrocytes ^73^, and in young actively remodeling bone ^74^, this, to our knowledge, is the first documented detection of Collagen Type VI in aged-human arthritic bone. This adds to a growing base of knowledge of the role of Collagen Type VI as a fibrotic biomarker outside its role in tissue development ^75^, now specifically in OA. The detection of various peptides of Collagen alpha-1(VI) and Collagen alpha-3(VI) (**Figure 5C, Supplemental Table 2**), may be the first detection of this network forming collagen in adult bone tissue outside the growth plate where chondrocytes aid in bone formation through endochondral ossification ^76,77^. Knockout of the *Col6a2* gene prevented normal ossification of the temporal mandibular joint in mice, suggesting that Collagen Type VI may play important roles in chondrocyte differentiation and bone growth in development ^76^, however its role in *bone* in OA is unknown. Collagen Type VI is produced by hypertrophic chondrocytes in arthritic cartilage where it can modulate pericellular mechanics ^73,78,79^. Strikingly, our detection of Collagen Type VI in OA subchondral bone hints at an important role for this collagen subtype in OA related bone remodeling and fibrosis. Although precise mechanisms behind this and the cell type responsible for its presence in osteoarthritic bone are currently unknown, the role of Collagen Type VI during bone development implies that this collagen subtype may mark the effort to mount a regenerative response, which may ultimately contribute to subchondral bone sclerosis in OA. While the peptide disease signature contains many different collagen peptides, the specific peptide identity and their functional and biological roles offer insight into the cellular and molecular mechanisms at play in subchondral bone in OA.

## Conclusions and Future Outlook

Overall, this study presents the first use of MALDI-MSI to investigate skeletal proteins in human joint disease providing further molecular knowledge about the progression of joint disease in subchondral bone. We show that markers of joint disease and OA are detectable in the subchondral bone, that these may be evident prior to cartilage loss, and that novel, specific regions of common ECM proteins may be hallmarks of this damage. Thus, by defining the spatial molecular landscape of OA on the protein level, this study advances new opportunities to increase the specificity of disease diagnostics and biomarkers for OA and opens new opportunities for future works with potential to establish new modes of disease tracking in human osteoarthritis. Further studies that implement MALDI-MSI to investigate the progression of human OA may be completed to extend the impact of this technological advance. First, there remain mainly promising molecular features and candidate peptides upregulated in our OA cohort that currently do not have matching IDs. In future work we will pursue these features by expanding existing skeletal tissue ECM specific libraries for the identification of novel biomarkers of disease. Additionally, emerging technologies are allowing *in situ* MS/MS (MS2) during MALDI-MSI that can accurately identify analytes during imaging experiments without the need to remove and digest imaged tissue. This will increase the ability to directly confirm protein IDs in place during imaging experiments, circumventing longer workflows utilizing LC-MS/MS for library building. Still, the promising success of this collagenase-based imaging motivates the use for other serial enzymatic digestions including elastases and chondroitinases ^44^ that can further probe the compositional changes of the ECM in OA. Excitingly, our new molecular evidence supports previous morphologic findings from microCT and MRI that subchondral bone in OA undergoes dramatic molecular remodeling, and these changes to subchondral bone can occur in bone even before cartilage loss in some cases ^5,6,10^. Thus, we suggest continued effort be spent in understanding the changes to subchondral bone as a way to develop diagnostics and treatments for osteoarthritis.

Osteoarthritis is a complex, personalized disease with multiple etiologies, with OA endotypes due to traumatic injury (post-traumatic OA), factors associated with obesity, and even normal healthy aging, among many others. Additionally, disease burden is skewed, such that cases of total knee replacement for woman are more than 33% higher than men^80^. Thus, we propose future studies utilizing MALDI-MSI enrolling larger cohorts of individuals with OA, both male and female, at multiple disease stages or ages and across several different endotypes of OA to assemble comprehensive knowledge of the molecular landscape of OA (**Figure 7**). The ability to visually detect and validate diagnostic biomarkers across ranges or subsets of patients would advance avenues of personalized medicine in the orthopedic field and in the context of OA. Paired with ongoing efforts to screen human synovial fluid for these markers, a route towards preemptive disease screening can be envisioned. Given the proteolytic environment of OA, naturally cleaved peptides are likely to exist in joint material and in the surrounding synovial fluid of inflamed OA joints, and these peptides may even make it to circulation. Thus, by identifying and tracking endogenous markers that increase with OA in subchondral bone, we can see a path towards the development of highly sensitive and specific diagnostics for a disease that currently severely lacks reliable early detection methods.

**Figure 7:**
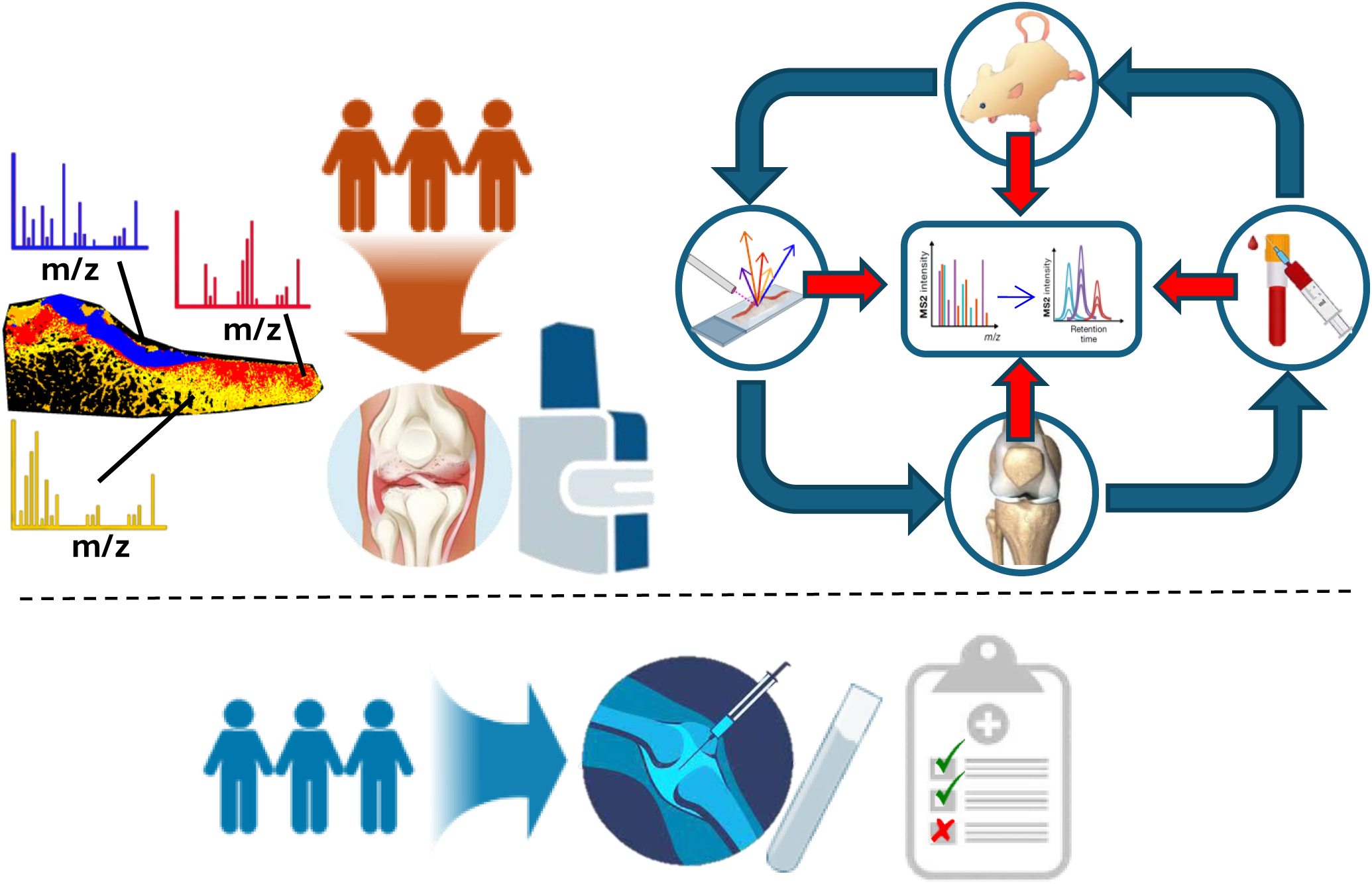
Potential Study Impact and Future Outlook. The study design for spatial Mass Spectrometry Imaging of OA patients potentially provided protein candidates that can be integrated within future studies to further explore OA mechanism and/or disease biomarkers. Future knee tissues from Total Knee Arthroplasty (TKA) patients from larger OA sample populations, including specific cohorts of male and female donors, will be used in similar foundational studies to expand the number of possible disease-specific protein and peptide features and biomarkers for OA. Validation studies will be performed *in vitro* and *in vivo*, as well as in OA mouse models. Validated OA biomarkers could (in the future) be used as potential diagnostic markers from synovial fluid or even circulating plasma to track OA risk and progression in a viable clinical method.

The extension of this technology to mouse models of OA where the biologic mechanism may be more precisely traced, and the disease progression and OA stage controlled, would provide much needed time course data and new hallmarks of early disease and tissue remodeling. Classification of the protein changes occurring in subchondral bone early in disease, prior to cartilage loss or the clinical presentation of pain, would greatly enhance the ability to develop diagnostics. In addition, knowledge gained from foundational mass spectrometry studies, such as this, can be used to identify precise molecular candidates from murine studies and larger panels of candidate peptides can be screened in biofluids from larger, healthy human populations (**Figure 7**). This circumvents the inherent difficulty and limitations of analyzing skeletal joint tissues from human patients with end-stage OA.

MALDI-MSI paired with targeted ECM enzymatic treatment has proven to be an effective tool to interrogate the molecular changes occurring in human joint tissue. We have demonstrated its potential for use in both discovery and potentially diagnostic based workflows, providing new much needed tools to investigate skeletal biology in a spatial manner. Findings from our work will motivate the use of MALDI-MSI and new technologies to further provide novel protein-based biomarkers for OA.

## Materials and Methods

### Human donor population and knee joint specimen preparation

Human knee joint tissues were obtained as described from the University of California, San Francisco, UCSF, as pre-prepared formalin-fixed paraffin-embedded (FFPE) tissue sections from an existing tissue bank ^7^. Briefly, four tibial plateaus from a group of non-OA diagnosed donors (cadaveric controls C1-C4) without a history of osteonecrosis, osteoporosis, or fractures were collected through the Willed Body Program at the University of California, San Francisco, and four osteoarthritic tibial plateaus were collected as surgical discard from patients receiving total knee arthroplasty/replacement, with an age range from 64 – 79 years old, (Patients P1-P4), who were clinically diagnosed with stage IV Osteoarthritis of the medial tibial plateau of the knee (**Supplemental Figures 1** and **2**). Recruitment occurred through referral from orthopedic surgeons at the Department of Veterans Affairs Medical Center (San Francisco, CA, USA). This resulted in an age, sex, and BMI matched cohort of human tibial plateaus that were stratified by their Osteoarthritis Research Society International (OARSI) score as graded by clinical pathologists as determined from histopathologic Safranin-O/Fast Green stains (**Supplemental Figures 1** and **2, Supplemental Table 1)**. All samples were collected with informed consent from patients with OA of the femorotibial joint as described in protocols that were reviewed and approved by the Human Subjects Protection Program Institutional Review Board of UCSF and the Department of Veterans Affairs Medical Center ^7^. All samples were harvested within 4 days postmortem to minimize the effects of degradation. Tibial plateaus, composed of the articular cartilage and underlying subchondral trabecular bone, were subsequently further divided into their medial and lateral halves through the posterolateral root of the posterior cruciate ligament, generating two independent tissue specimens for each patient tissue representing the lateral and medial halves of the same joint from a single patient to accommodate placement on standard 25×75 mm microscopy slides. This allowed for the comparison of medial OA sections (P1-P4, medial) to two distinct sets of ‘control’ tissues composed of i) site-matched non-arthritic cadaveric controls (C1-C4, medial) and ii) tissue-matched OA lateral tibial plateau halves (from the same OA patients, P1-P4, lateral). Lateral cadaveric control tissues were collected, processed, and imaged but resemble the cadaveric medial tissues in most cases.

### Tissue Preparation and Sectioning

Human tibial plateaus were fixed in 10% neutral buffered formalin (NBF) and incubated in an Ion Exchange Decalcification Unit (American Master Technologies) for 5–6 days until fully demineralized. Serial ethanol dehydration and paraffin infiltration and embedding were performed to generate FFPE tissue blocks. Tibial plateaus were sectioned onto standard pathology microscopy slides at 7 μm thickness in the coronal plane. Histological Safranin-O / Fast Green stain **(Supplemental Figure 2)** was used to identify regions of cartilage and cartilage damage in tissue sections. Unstained neighboring tissue sections were selected for Matrix-Assisted Laser Desorption Ionization – Mass Spectrometry Imaging (MALDI-MSI) preparation and imaging.

### Sample Preparation for MALDI-MSI for ECM Imaging

Formalin-fixed paraffin-embedded human tibial plateau tissue slides were prepared for enzymatic digestion using previously published protocols ^23,44,46,81^ with important modifications in consideration of the large, ECM-rich tissue sections of the human tibial plateaus. The slides were heated at 60 °C for 1 hour to melt off excess paraffin wax. After cooling, tissue sections were further deparaffinized by washing twice in xylene (3 minutes each). Tissue sections were washed and rehydrated by submerging in 100% ethanol twice (1 minute each), once in a Carnoy’s solution (60% ethanol, 30% chloroform and 10% glacial acetic acid), to remove excess fat and wax, once in 95% ethanol (one minute), once in 70% ethanol (one minute), and twice in water (3 minutes each). After the rehydration series, slides were transferred to a slide mailer containing citraconic anhydride buffer (Thermo) at pH 3.0 for heat induced antigen/epitope retrieval. The slide mailer was sealed and gently arranged in a 60 °C oven so that submerged slides were positioned face up while the mailers were additionally submerged in a larger container (glass beaker) containing buffer that allowed for their orientation and provided an additional buffer reservoir, and left to incubate overnight, not exceeding 12 hours. Slide mailers were then removed from the oven and allowed to cool on the bench top. Citraconic buffer was slowly exchanged with water, as to not impact the tissue, five times by removing half of the buffer via pipette and replacing it with water, prior to replacing it completely with water for the final solution exchange. Deglycosylation was achieved by applying a 0.1 μg/μL PNGase F solution (N-Zyme Scientifics) via the TM Sprayer M3 (HTX Imaging, Chapel Hill, NC, USA), followed by incubation in custom humid chambers for 2 hours at 37°C. After deglycosylation, slides were coated with a 7 mg/mL solution of alpha-cyano-4-hydroxycinnamic acid (CHCA) 50% acetonitrile, 0.1% trifluoracetic acid matrix solution via the TM Sprayer M3. After glycan imaging or deglycosylation, slides were washed carefully with 70% ethanol to remove the applied CHCA matrix, a high pH buffer (10 mM Tris, pH 9, and low pH buffer, 10 mM citraconic anhydride, pH 3, followed by distilled water to remove glycans and residual enzyme. Specific care was taken to ensure that serial enzymatic digestion post glycan imaging would not disrupt the integrity of the tissue slides ^44,45^. Following the wash, slides were vacuum dried in a desiccator and prepared for ECM peptide imaging performing a second epitope retrieval (10 mM Tris, pH 9 overnight at 60 °C). ECM-targeted enzymatic digestion for proteomic imaging was completed utilizing collagenase III that featured an enzymatic preference to cleave at repeated amino acid Gly-Xxx-Pro (GXP) motifs within the triple helical region of collagens ^82,83^. While preferably targeting collagens, this enzyme formulation also allowed for the detection of several other ECM and ECM-related proteins (**Supplemental Table 2**). The proteolytic enzyme pre-treatment was composed of a 0.1 μg/μL collagenase III (COLase3 (*Clostridium histolyticum*) Worthington Biochemical, Lakewood, NJ, USA) solution applied via the TM Sprayer M3, and incubated for 5 hours in humid chambers at 37 °C. After incubation, slides were vacuum dried in a desiccator and coated with CHCA matrix (7 mg/mL solution of alpha-cyano-4-hydroxycinnamic acid (CHCA) 50% acetonitrile, 1.0% trifluoracetic acid). The CHCA matrix solution was spiked with [Glu1]-fibrinopeptide B human (GluFib) (Sigma-Aldrich, St Louis, MO, USA) for a total concentration of 200 fmol/L. After matrix drying for 30 minutes in a desiccator, the slides were rapidly immersed in cold 5 mM ammonium phosphate, monobasic for <1 second and immediately dried in a desiccator.

### MS Imaging Data Processing and Statistical Analysis

Image processing and mass spectrometry data analysis were completed in SCiLS Lab MVS v2024a Pro (Bruker Scientific, LLC, Bremen, Germany). During import, SCiLS lab background normalizes and resamples the raw data to ensure a common mass axis across the several thousand independent spectra acquisitions using a TIC preserving algorithm. This is accomplished through the SCiLS lab Import Wizard ^84^. After importation, each composite spectrum is further normalized to the glufibrinogen (GluFib) standard peptide at m/z 1570.6768 such that the remaining spectra have the same ion count in the peak interval (unit area under the peak) and the rest of each spectrum is scaled accordingly^85^. Backgrounded and normalized spectra are additionally filtered such that only the top 500 most abundant m/z features from each tissue are used in subsequent data analysis to avoid residual noise. These selected 500 peaks are automatically detected and used to generate tissue regions through a Bisecting K-means method using the Segmentation operation within SCiLS Lab ^86^. This unbiased segmentation recognized regions of tissue that shared similar spectral profiles and grouped them together into regions. The number of hierarchical regions generated was set to the minimum required to identify cartilage regions from bone and to resolve subchondral trabecular bone against the background or marrow space areas. This unsupervised processing typically generated 4 to 6 tissue subregions often identifying multiple transitional bone subtypes. These generated regions are used to perform paired Discriminating Feature analysis ^51,87,88^ within SCiLS Lab. These analyses processed all 30,000-40,000 captured spectra per imaged tissue (specific number of spectra differed by tissue size per sample) to identify candidate marker features (m/z) defining the difference between identified non-arthritic and diseased subchondral bone regions in both site-matched (cadaveric “healthy donor medial to OA medial tissues) and tissue-matched (OA patient lateral vs OA patient medial) comparisons. Candidate markers were considered if the area under the receiver operating curve (ROC) generated by the Discriminating Feature analysis for a given m/z was ≥0.85. Candidates were first generated by comparing one representative sample from the control group to the OA group. Candidates were later validated across the whole cohort by exporting intensity values for each candidate m/z out of SCiLS lab. Technical and computational limitations on file size prevented simultaneous analysis of numerous large files at once in SCiLS lab. To ensure then that candidate lists were not overly sample specific, similar sets of candidate peptides were generated using tissue sections from different individuals (**Supplemental Figure 7**). In addition to the automated tissue segments from SCiLS Lab, histologic stains of serial tissue sections were used to draw regions of interest (ROIs) manually. These ROIs included the diseased regions of subchondral bone to the edge of exposed cartilage on the medial side in OA patients or an equal distance of subchondral bone of the lateral side of the tissue matched joint (**Figure 5A**), or ∼2/5 the length of subchondral bone in cadaveric controls from the medial edge towards the center (the approximate distance of lost cartilage in medial OA samples) (**Supplemental Figure 5**). The individual pixel intensities by ROI for each candidate m/z feature were exported from SCiLS and the mean pixel intensity was calculated for each candidate feature per each ROI (patient or control sample). Peptide identifications were assigned to m/z features by searching the measured MS1 singly charged precursor ions (M+H) and their m/z value for each feature against previously generated peptide libraries assembled from other tissue types, including breast, cardiac, and lung tissues ^20,42,46^. Identifications were considered valid if the mass difference between the MALDI feature m/z and the peptide library were ≤ 2 mDa (∼15 ppm) from the detected m/z value on the MS1 level in the MALDI-MSI to matching values in the separate databases that had been confirmed in greater detail in LC-MS/MS. In some cases, detected m/z features had multiple potential peptide or protein matches in databases and so the multiple ID sequences are reported (**Supplemental Table 2**), but overall statistical analysis considers features with multiple matches as a single candidate feature for reporting purposes. Further MS/MS is needed to confirm tissue-specific peptide sequence matches. In Results, only features with unique identifications are presented and discussed. MetaboAnalyst v5.0 ^89^ was used to generate three dimensional partial least squares-discriminant analysis (PLS-DA) plots using mean feature pixel intensity from the combined set of unique candidate features obtained through ROC analysis for each group comparison type. Although some candidate features were originally found from ROC analysis in one comparison or another (cadaveric Medial vs OA Medial or OA Medial vs OA Lateral), they exist as detectable features in all samples and statistical testing for group-wise comparisons between OA medial and the two control groups utilized the full feature list. Candidate m/z features were “validated” in group-wise comparisons using the sample mean intensity for individual candidate features from each independent tissue sample with Prism Version 10.1.1 (GraphPad) utilizing a paired or unpaired one-sided student’s t-test as appropriate (e.g., within patient tissue or between different individuals) with significance set *a priori* at p < 0.05. A one-sided t-test was selected given that intensity distributions from each ROI were right-skewed and intensity values cannot be negative values. Biological replicates of N=4 for all groups.

### Mass Spectrometry Imaging

Mass spectrometric imaging data after proteolytic digestion with collagenase III was acquired using a Trapped Ion-Mobility Time-of-Flight Mass Spectrometer, the timsTOF fleX (Bruker Scientific, LLC, Bremen, Germany) equipped with a dual ESI/MALDI source. Samples were scanned across a MS1 mass range at m/z 600 – 4000. The SmartBeam 3D 10 kHz laser was set to 20% power, scan range of 20 μm for X and Y and resulting field size of 24 μm for X and Y. 300 spectra were collected per pixel spot. Additional instrument parameters for the studies included: an ion transfer time of 75.0 μs, pre pulse storage time of 20 μs, a collision RF of 2500 Vpp, a collision energy of 25 eV, an ion energy in the quadrupole of 5.0 eV, a TIMS funnel 1 RF of 400 Vpp, a TIMS funnel 2 RF of 500 Vpp and a multipole RF of 400 Vpp. After MS acquisition, the data was imported into SCiLS Lab MVS v2024a Pro (Bruker Scientific, LLC, Bremen, Germany). MSI generates spatial maps of the imaged tissue in which the intensities correspond to the abundance of a molecular feature (e.g., peptide) at the imaged location with a spot size of 20 μM, with variable step-size resolution from adjacent pixels at 20 μM laser step size to capture small regions in detail to a larger step sizes of 120 μM or more to create images of the entire tissue section..

### Tissue Preparation and Electrospray Ionization MS/MS by Data-Dependent Acquisition

After imaging, the MALDI matrix was removed by 70% ethanol wash. Tissue was removed from slides and fully digested in solution with the same collagenase III used to prepare slides for imaging. Excess enzyme removal and desalting of digested peptides were completed by sequential stage tip and ZipTip (Merk Millipore, Burlington MA) purification. Samples (100 ng) were loaded onto EvoTips Pure (Evosep, Odense, Denmark) following the manufacturer’s protocol. LC-MS/MS analyses were performed on an Evosep One liquid chromatography system (Evosep) coupled to a timsTOF HT mass spectrometer (Bruker, Bremen, Germany). The solvent system consisted of 0.1% formic acid (FA) in water (solvent A) and 0.1% FA in ACN (solvent B). Peptides were eluted on a PepSep C_18_ analytical column (150 µm x 15 cm, 1.5 µm particle size; Bruker) using the 30 SPD method (44-minute gradient length, 500-nL/min flow rate). A zero-dead volume emitter was installed in the nano-electrospray source (CaptiveSpray source, Bruker Daltonics) and the source parameters were set as follows: capillary voltage 1600 V, dry gas 3 L/min, and dry temperature 180°C. Each sample was acquired in data-dependent acquisition (DDA) - parallel accumulation–serial fragmentation (PASEF) ^90^ mode with 4 PASEF MS/MS ramps and with the trapped-ion mobility (TIMS) mobilogram inclusion parallelogram expanded to include monocharged precursor ions (**Supplemental Table 3**). For DDA-PASEF with TIMS on, MS1 and MS2 spectra were acquired over an m/z range of 100-1700 m/z and an ion mobility range of 0.85 -1.70 Vs/cm^2^. Precursor ions were selected fragmentation with charge state from 0 to 5, target intensity of 12,500, and intensity threshold of 500. The dual TIMS analyzer was operated in a 100% duty cycle with equal accumulation time and ramp time of 150 ms each, for a total cycle time of 0.78 s. The collision energy was defined as a linear function of mobility starting from 20 eV at 1/K0 = 0.6 Vs/cm^2^ to 59 eV at 1/K0 = 1.6 Vs/cm^2^. For calibration of ion mobility dimension, three ions of Agilent ESI-Low Tuning Mix ions were selected (m/z [Th], 1/K0 [Vs/cm^2^]: 622.0289, 0.9915; 922.0097, 1.1986; 1221.9906, 1.13934). Spectral libraries from acquisitions were generated utilizing Spectromine version 4.0 (Biognosys AG, Schlieren, Switzerland) using a nonspecific enzymatic digestion with peptide length of 7-52 amino acids, with fixed cysteine carbamidomethyl modifications and variable N-terminal protein acetylation and oxidation of methionine and proline. Protein, peptide, and PSM FDR were set to 0.01. Searches were made against a Human UniProtKB/SwissProt reviewed database downloaded from UniProt on April 3, 2023, containing 1032 entries of proteins with “Extracellular Matrix” GO cellular component annotation. Finally, the best 3-6 fragments per peptide were kept. MS/MS spectra for hydroxyproline containing peptides were manually inspected and verified in Skyline-daily v21.2.1.535 (University of Washington, WA, USA) ^91^. The final library contained 543 peptides, 320 of which contained hydroxyproline modifications, all corresponding to 58 protein groups, all spectra were manually reviewed. For peptides where DDA analysis was unable to specifically determine HYP modification site location due to multiple potential prolines within a peptide, **Supplemental Table 2** lists the PTM site score probabilities as determined by MaxQuant for each modified peptide as reported by other databases^92^.

## Data Availability

The human spatial proteomic datasets have been uploaded to the Center for Computational Mass Spectrometry MassIVE repository at UCSD and can be accessed using the following link: https://massive.ucsd.edu/ProteoSAFe/private-dataset.jsp?task=c8ee2b4247994c4396007e1cffd0bcfb

**[Note to reviewers]:** To access data prior during revision (before public release) please utilize the following information: MassIVE ID number: **MSV000094448**; Username: **MSV000094448_reviewer**, Password: winter.

## Acknowledgements

We acknowledge the NIH for support from the following granting agencies and mechanisms: NIA T32 AG000266 (to Schurman, PI: Ellerby), NIA P01 AG066591 – (PI: Ellerby, Core Director: Schilling), NIA U01 AG060906 – (PI: Schilling), NIDCR R01 DE019284 – (PI: Alliston), DOD PRORP W81XWH1810155 (PI: Alliston), NIAMS P30 AR066262 (PI: Alliston), DOD PRORP OR130191 (PI: Alliston), NIAMS R21 AR083065 (PI: Alliston), DOD HT94252310875 (PI: Alliston), NCI R21 CA240148 (PI: Angel), NIAMS R21 AR084303 (MPIs: Schilling, Alliston, Angel) and NIGMS/OD S10 OD030212 (PI: Angel). Drs. Alfred Kuo and Thomas Vail supported the study with the experimental design, collection, and original analysis of human tissue samples. Drs. Schurman and Schilling like to recognize the amazing support and memory of Dr. Judith Campisi for her lifelong dedication to science and advocation for scientific trainees, especially in the field of aging biology.

---

**Supplementary Figure 1: Male OA Patient and Control Cohort Data:** (A) Fresh control male cadaveric tissues (n=4) were obtained through the Willed Body Program at University of California, San Francisco from age, sex, and BMI matched donors without history of OA, osteonecrosis, osteoporosis, or fractures. Tibial Plateaus from four 65-74 yr. old male subjects with clinically diagnosed stage IV osteoarthritis (OA) receiving total knee replacement were recruited through referral from orthopedic surgeons at the Department of Veterans Affairs Medical Center, San Francisco. * p < 0.05 in by student’s T-Test showed that the control and OA patient cohorts were distinguished by their OARSI grade (B) but were both BMI (C) and age (D) matched.

**Supplemental Figure 2: OA & Control Tissue Sections and Safranin O Staining.** (A) Unstained tissue scans and (B) Safranin-O Histologic staining of FFPE slides of medial and lateral halves of the Tibial Plateau of TKA Patients and the lateral half from donor cadaveric non-arthritic Controls. Safranin O staining appears as red to stain proteoglycans in cartilage layers while the subchondral bone remains white/grey.

**Supplemental Figure 3: MALDI-MSI Identified Molecular Features that Define the Skeletal ECM.** (A) Isolation of specific m/z features from the composite spectrum of all imaged pixels allowed for the generation of spatially resolved heatmaps of peptide distribution. Isolation of individual peaks in the composite spectrum, of over 3,000 individual peaks, revealed peptide ions that were preferentially found in bone or cartilage regions, others were detected across both tissues. (B) The top 500 most abundant peaks from the composite spectrum were used to perform automated region segmentation using a bisecting K-means method. Although only using data from on a subset of peaks from the full composite spectrum, this segmentation operation accurately captured 3 regions in non-arthritic, aged tissue that physiologically matched i) the cartilage (blue), ii) the subchondral bone (orange), iii) and bone marrow (maroon) regions between trabecula; additional ‘background’ or pixels with little tissue specific signal (black) were also identified. (C) The hierarchical segmentation tree showed how many pixels are assigned into each of the generated regions based on their independent spectra.

**Supplemental Figure 4: Additional Spatial Markers by MALDI-MSI of Skeletal Tissue.** (A) Additional unique molecular features or peptides of skeletal marker proteins Collagen alpha-1(I), G^626^PAGERGEQ^634^ at m/z 900.40, for bone, and Collagen alpha-1(II), G^998^PSGEPGKQGAP^1010^, for cartilage at m/z 1097.53, were preferentially detected in the identified bone and cartilage areas demonstrating the agreement between the automatically generated regions and known skeletal tissues. *p < 0.05 of average pixel intensity in bone (orange) vs cartilage (blue). (B) High resolution scanning (20-micron step) in a tissue region composed of the bone and cartilage junction showed the spatial delineation of tissue type and spatial distribution of peptides within trabecular bone. Interestingly this peptide of Collagen alpha-1(II) shows faint signal within trabecula but was most prominently contained with the cartilage region. High resolution images of the subchondral bone / cartilage junction for unidentified peptide features at (C) m/z 1943.91, (D) m/z 1406.71, and (E) m/z 1242.58 showed additional unique spatial distributions for different molecular features and ‘hot spots’ within the subchondral trabecular bone with low signal from the cartilage region.

**Supplemental Figure 5:** OA Medial vs Cadaveric Medial Comparison. (A) Site-matched comparison of arthritic OA medial tibial plateaus to age and BMI matched non-arthritic cadaveric tibial plateaus. Outlined blue areas show the identified cartilage tissue in OA and Cadaveric tissue used to delineate the subchondral bone regions of interest (ROI) (Green) used for quantitative analysis. (B) Three-dimensional PLS-DA of candidate feature intensity from the segmented ROIs confirmed group separation in multiple component dimensions.

**Supplemental Figure 6: Gradient of Disease in Age-related Osteoarthritis.** A) Heatmap for a candidate peptide, GLP^(ox)^GERGRP^(ox)^GAP^(ox)^ at m/z 1211.61, from Collagen alpha-1(I) (Col1A1) that was found upregulated in OA is depicted across both halves of Cadaveric control C1 and OA patient P1 tissue sections. B) Quantification of normalized pixel intensity for the same peptide showed gradient of increase from Cadaveric Lateral to OA Medial tissue sections. All four samples were statistically different from one another by post-hoc Tukey testing after one-way ANOVA (p<0.05). Inset (dotted box) shows increased resolution on the y-axis (between 0 and 0.15) around the mean pixel intensity values for each tissue section to more precisely depict differences in feature abundance across the four tissue sections.

**Supplemental Figure 7: Tissue Segmentation and candidate ion generation from additional OA Patients.** OA Patient 3 (P3) (A) was used to generate candidate features. (B) The top 500 most abundant m/z features were used to automatically generate tissue regions. (C) Extraction of the tissue regions containing sclerotic diseased bone and non-arthritic region allowed for feature analysis. (D) ROC analysis between the extracted regions resulted in 37 candidate ions that described the difference between the independent regions with an AUROC of > 0.85. (E) Comparison of the 37 specific ions identified from patient #3 shared 19 ions in common with candidates generated from those identified using representative patient #1.

**Supplemental Figure 8: Tissue-matched OA Medial vs Lateral Candidate m/z Comparisons across entire cohort.** Intensities for 30 of 44 candidates are significantly increased in OA across the patient cohort by paired one-sided t-test (p< 0.05). 8 additional markers trended towards significance (0.1 < p < 0.05).

**Supplemental Figure 9**: **Site-matched OA Medial vs CAD Medial Candidate m/z Comparisons across entire cohort.** Intensities for 7 of 44 candidates significantly increased in OA across the patient cohort by one-sided t-test. (p<0.05) 10 additional markers trended towards significance (0.1 < p < 0.05).

**Supplemental Figure 10: MS/MS Spectra for Hydroxyproline-Modified Peptides.** Data-Dependent Acquisition (DDA) MS/MS spectra of ions corresponding to MALDI-MSI features from Figure 6 matching peptides originating from Collagen alpha-1(I) (Col1A1) profiled with 1 or more modified hydroxyproline residues. Fragment ions from each precursor ion confirm the presence of modified hydroxyproline residues. (A) A singly modified peptide I^954^AGQRGVVGLP^(ox)964^, at m/z 1082.63 was detected with multiple y-ions containing the hydroxyproline residue from this precursor. (B) MS/MS spectra displaying fragment ions of the peptide G^656^KP^(ox)^GEQGVP^(ox)^GDLGAP^(ox)670^ at m/z 1426.68 containing 2 HYP residues showed a mass shift in b- and y-ions at m/z by 16 m/z (15.994) at b_9_ and y_7_ ions confirming the extra HYP modification. Note that in (C) fragment ions b_6_ and y ^++^ shared the same m/z in the MS/MS scans.

**Supplemental Figure 11: Bioactive Regions on Collagen Type 1 Upregulated in OA.** A) The Collagen alpha-1(I) (Col1A1) peptide G^1165^PRGRTGD^1173^ at m/z 1139.58 is located in a region of the mature collagen fibril with multiple binding partners including, Amyloid Precursor Protein, Interleukin 2, and Phosphophoryn. B) Another Collagen alpha-1(I) peptide upregulated in medial OA tissues, G^626^PAGERGEQ^634^ overlaps a known glycation site and a proposed region for keratan sulfate interaction. Protein sequences of the mature Collagen 1 triple helix are shown as GenBankTM, 121(I) accession #NP000079.2 and 122(I) NP_000080.2. Amino Acid positions noted are from UniProt P02452 CO1A1_HUMAN and P08123 CO1A2_HUMAN. Collagen fibril binding zones schematics adapted from “Antonio, J. D. S., Jacenko, O., Fertala, A. & Orgel, J. P. R. O. Collagen Structure-Function Mapping Informs Applications for Regenerative Medicine. *Bioeng* 8, 3 (2020)”. ^1^PHOS: Phosphophoryn binding site. ^2^C-Pep. Domain: C-telopeptide fibrillogenesis nucleation domain. ^3^KSPG: KSPG - Proposed sites of keratan sulfate proteoglycan binding.

## References

1 Fuchs, J., Kuhnert, R. & Scheidt-Nave, C. 12-month prevalence of osteoarthritis in Germany. J Health Monit 2, 51–56 (2017). 10.17886/RKI-GBE-2017-066

2 Resende, V. A. C. et al. Higher age, female gender, osteoarthritis and blood transfusion protect against periprosthetic joint infection in total hip or knee arthroplasties: a systematic review and meta-analysis. Knee Surg Sports Traumatol Arthrosc 29, 8–43 (2021). 10.1007/s00167-018-5231-9

3 Schurman, C. A. et al. Molecular and Cellular Crosstalk between Bone and Brain: Accessing Bidirectional Neural and Musculoskeletal Signaling during Aging and Disease. J Bone Metab 30, 1–29 (2023). 10.11005/jbm.2023.30.1.1

4 Heidari, B. Knee osteoarthritis prevalence, risk factors, pathogenesis and features: Part I. Caspian J Intern Med 2, 205–212 (2011).

5 Burr, D. B. & Gallant, M. A. Bone remodelling in osteoarthritis. Nat Rev Rheumatol 8, 665–673 (2012). 10.1038/nrrheum.2012.130

6 Goldring, S. R. & Goldring, M. B. Changes in the osteochondral unit during osteoarthritis: structure, function and cartilage-bone crosstalk. Nat Rev Rheumatol 12, 632–644 (2016). 10.1038/nrrheum.2016.148

7 Mazur, C. M. et al. Osteocyte dysfunction promotes osteoarthritis through MMP13-dependent suppression of subchondral bone homeostasis. Bone Res 7, 34 (2019). 10.1038/s41413-019-0070-y

8 Hu, Y., Chen, X., Wang, S., Jing, Y. & Su, J. Subchondral bone microenvironment in osteoarthritis and pain. Bone Res 9, 20 (2021). 10.1038/s41413-021-00147-z

9 Alliston, T., Hernandez, C. J., Findlay, D. M., Felson, D. T. & Kennedy, O. D. Bone marrow lesions in osteoarthritis: What lies beneath. J Orthop Res 36, 1818–1825 (2018). 10.1002/jor.23844

10 Namiri, N. K. et al. Deep learning for large scale MRI-based morphological phenotyping of osteoarthritis. Sci Rep 11, 10915 (2021). 10.1038/s41598-021-90292-6

11 Hayami, T. et al. The role of subchondral bone remodeling in osteoarthritis: reduction of cartilage degeneration and prevention of osteophyte formation by alendronate in the rat anterior cruciate ligament transection model. Arthritis Rheum 50, 1193–1206 (2004). 10.1002/art.20124

12 Cabral, W. A. et al. Prolyl 3-hydroxylase 1 deficiency causes a recessive metabolic bone disorder resembling lethal/severe osteogenesis imperfecta. Nat Genet 39, 359–365 (2007). 10.1038/ng1968

13 Saito, M. & Marumo, K. Effects of Collagen Crosslinking on Bone Material Properties in Health and Disease. Calcif Tissue Int 97, 242–261 (2015). 10.1007/s00223-015-9985-5

14 Acevedo, C. et al. Contributions of Material Properties and Structure to Increased Bone Fragility for a Given Bone Mass in the UCD-T2DM Rat Model of Type 2 Diabetes. J Bone Miner Res 33, 1066–1075 (2018). 10.1002/jbmr.3393

15 Schurman, C. A. et al. Aging impairs the osteocytic regulation of collagen integrity and bone quality. Bone Res 12, 13 (2024). 10.1038/s41413-023-00303-7

16 Franck, J. et al. MALDI imaging mass spectrometry: state of the art technology in clinical proteomics. Mol Cell Proteomics 8, 2023–2033 (2009). 10.1074/mcp.R800016-MCP200

17 Walch, A., Rauser, S., Deininger, S. O. & Hofler, H. MALDI imaging mass spectrometry for direct tissue analysis: a new frontier for molecular histology. Histochem Cell Biol 130, 421–434 (2008). 10.1007/s00418-008-0469-9

18 Michno, W., Wehrli, P. M., Blennow, K., Zetterberg, H. & Hanrieder, J. Molecular imaging mass spectrometry for probing protein dynamics in neurodegenerative disease pathology. J Neurochem 151, 488–506 (2019). 10.1111/jnc.14559

19 Hanrieder, J., Ljungdahl, A. & Andersson, M. MALDI imaging mass spectrometry of neuropeptides in Parkinson’s disease. J Vis Exp (2012). 10.3791/3445

20 Angel, P. M. et al. Extracellular Matrix Imaging of Breast Tissue Pathologies by MALDI-Imaging Mass Spectrometry. Proteomics Clin Appl 13, e1700152 (2019). 10.1002/prca.201700152

21 Angel, P. M. et al. Zonal regulation of collagen-type proteins and posttranslational modifications in prostatic benign and cancer tissues by imaging mass spectrometry. Prostate 80, 1071–1086 (2020). 10.1002/pros.24031

22 Heijs, B. et al. Molecular signatures of tumor progression in myxoid liposarcoma identified by N-glycan mass spectrometry imaging. Lab Invest 100, 1252–1261 (2020). 10.1038/s41374-020-0435-2

23 Angel, P. M., Mehta, A., Norris-Caneda, K. & Drake, R. R. MALDI Imaging Mass Spectrometry of N-glycans and Tryptic Peptides from the Same Formalin-Fixed, Paraffin-Embedded Tissue Section. Methods Mol Biol 1788, 225–241 (2018). 10.1007/7651_2017_81

24 Tuck, M., Grelard, F., Blanc, L. & Desbenoit, N. MALDI-MSI Towards Multimodal Imaging: Challenges and Perspectives. Front Chem 10, 904688 (2022). 10.3389/fchem.2022.904688

25 Ogrinc, N. et al. Robot-Assisted SpiderMass for In Vivo Real-Time Topography Mass Spectrometry Imaging. Anal Chem 93, 14383–14391 (2021). 10.1021/acs.analchem.1c01692

26 Eveque-Mourroux, M. R., Rocha, B., Barre, F. P. Y., Heeren, R. M. A. & Cillero-Pastor, B. Spatially resolved proteomics in osteoarthritis: State of the art and new perspectives. J Proteomics 215, 103637 (2020). 10.1016/j.jprot.2020.103637

27 Aichler, M. & Walch, A. MALDI Imaging mass spectrometry: current frontiers and perspectives in pathology research and practice. Lab Invest 95, 422–431 (2015). 10.1038/labinvest.2014.156

28 Kriegsmann, M. et al. MALDI MS imaging as a powerful tool for investigating synovial tissue. Scand J Rheumatol 41, 305–309 (2012). 10.3109/03009742.2011.647925

29 Cillero-Pastor, B., Eijkel, G. B., Blanco, F. J. & Heeren, R. M. A. Protein classification and distribution in osteoarthritic human synovial tissue by matrix-assisted laser desorption ionization mass spectrometry imaging. Analytical and Bioanalytical Chemistry 407, 2213–2222 (2015). 10.1007/s00216-014-8342-2

30 Lee, Y. R. et al. Mass Spectrometry Imaging as a Potential Tool to Investigate Human Osteoarthritis at the Tissue Level. Int J Mol Sci 21 (2020). 10.3390/ijms21176414

31 Rocha, B. et al. Identification of a distinct lipidomic profile in the osteoarthritic synovial membrane by mass spectrometry imaging. Osteoarthritis Cartilage 29, 750–761 (2021). 10.1016/j.joca.2020.12.025

32 Eveque-Mourroux, M. R. et al. Spatially resolved endogenous improved metabolite detection in human osteoarthritis cartilage by matrix assisted laser desorption ionization mass spectrometry imaging. Analyst 144, 5953–5958 (2019). 10.1039/c9an00944b

33 Haartmans, M. J. J. et al. Matrix-assisted laser desorption/ionization mass spectrometry imaging (MALDI-MSI) reveals potential lipid markers between infrapatellar fat pad biopsies of osteoarthritis and cartilage defect patients. Analytical and Bioanalytical Chemistry 415, 5997–6007 (2023). 10.1007/s00216-023-04871-9

34 Schaepe, K. et al. Imaging of Lipids in Native Human Bone Sections Using TOF-Secondary Ion Mass Spectrometry, Atmospheric Pressure Scanning Microprobe Matrix-Assisted Laser Desorption/Ionization Orbitrap Mass Spectrometry, and Orbitrap-Secondary Ion Mass Spectrometry. Anal Chem 90, 8856–8864 (2018). 10.1021/acs.analchem.8b00892

35 Seeley, E. H. et al. Co-registration of multi-modality imaging allows for comprehensive analysis of tumor-induced bone disease. Bone 61, 208–216 (2014). 10.1016/j.bone.2014.01.017

36 Fujino, Y., Minamizaki, T., Yoshioka, H., Okada, M. & Yoshiko, Y. Imaging and mapping of mouse bone using MALDI-imaging mass spectrometry. Bone Rep 5, 280–285 (2016). 10.1016/j.bonr.2016.09.004

37 Svirkova, A., Turyanskaya, A., Perneczky, L., Streli, C. & Marchetti-Deschmann, M. Multimodal imaging of undecalcified tissue sections by MALDI MS and muXRF. Analyst 143, 2587–2595 (2018). 10.1039/c8an00313k

38 Lee, Y. R. et al. High-Resolution N-Glycan MALDI Mass Spectrometry Imaging of Subchondral Bone Tissue Microarrays in Patients with Knee Osteoarthritis. Anal Chem 95, 12640–12647 (2023). 10.1021/acs.analchem.3c00348

39 Briggs, M. T. et al. MALDI mass spectrometry imaging of N-glycans on tibial cartilage and subchondral bone proteins in knee osteoarthritis. Proteomics 16, 1736–1741 (2016). 10.1002/pmic.201500461

40 Cillero-Pastor, B., Eijkel, G. B., Kiss, A., Blanco, F. J. & Heeren, R. M. Matrix-assisted laser desorption ionization-imaging mass spectrometry: a new methodology to study human osteoarthritic cartilage. Arthritis Rheum 65, 710–720 (2013). 10.1002/art.37799

41 Rocha, B., Cillero-Pastor, B., Blanco, F. J. & Ruiz-Romero, C. MALDI mass spectrometry imaging in rheumatic diseases. Biochim Biophys Acta Proteins Proteom 1865, 784–794 (2017). 10.1016/j.bbapap.2016.10.004

42 Angel, P. M. et al. Extracellular matrix alterations in low-grade lung adenocarcinoma compared with normal lung tissue by imaging mass spectrometry. J Mass Spectrom 55, e4450 (2020). 10.1002/jms.4450

43 Clift, C. L. et al. Evaluation of Therapeutic Collagen-Based Biomaterials in the Infarcted Mouse Heart by Extracellular Matrix Targeted MALDI Imaging Mass Spectrometry. J Am Soc Mass Spectrom 32, 2746–2754 (2021). 10.1021/jasms.1c00189

44 Clift, C. L., Drake, R. R., Mehta, A. & Angel, P. M. Multiplexed imaging mass spectrometry of the extracellular matrix using serial enzyme digests from formalin-fixed paraffin-embedded tissue sections. Anal Bioanal Chem 413, 2709–2719 (2021). 10.1007/s00216-020-03047-z

45 Clift, C. L., Mehta, A., Drake, R. R. & Angel, P. M. Multiplexed Imaging Mass Spectrometry of Histological Staining, N-Glycan and Extracellular Matrix from One Tissue Section: A Tool for Fibrosis Research. Methods Mol Biol 2350, 313–329 (2021). 10.1007/978-1-0716-1593-5_20

46 Angel, P. M. et al. Advances in MALDI imaging mass spectrometry of proteins in cardiac tissue, including the heart valve. Biochim Biophys Acta Proteins Proteom 1865, 927–935 (2017). 10.1016/j.bbapap.2017.03.009

47 Babatunde, O. A. et al. Racial Distribution of Neighborhood-Level Social Deprivation in a Retrospective Cohort of Prostate Cancer Survivors. Diseases 10 (2022). 10.3390/diseases10040075

48 Oegema, T. R., Jr., Carpenter, R. J., Hofmeister, F. & Thompson, R. C., Jr. The interaction of the zone of calcified cartilage and subchondral bone in osteoarthritis. Microsc Res Tech 37, 324–332 (1997). 10.1002/(SICI)1097-0029(19970515)37:4<324::AID-JEMT7>3.0.CO;2-K

49 Sandhu, A., Rockel, J. S., Lively, S. & Kapoor, M. Emerging molecular biomarkers in osteoarthritis pathology. Ther Adv Musculoskelet Dis 15, 1759720X231177116 (2023). 10.1177/1759720X231177116

50 Cook, N. R. Statistical evaluation of prognostic versus diagnostic models: beyond the ROC curve. Clin Chem 54, 17–23 (2008). 10.1373/clinchem.2007.096529

51 Hoo, Z. H., Candlish, J. & Teare, D. What is an ROC curve? Emerg Med J 34, 357–359 (2017). 10.1136/emermed-2017-206735

52 Greene, M. A. & Loeser, R. F. Aging-related inflammation in osteoarthritis. Osteoarthritis Cartilage 23, 1966–1971 (2015). 10.1016/j.joca.2015.01.008

53 Li, Y., Wei, X., Zhou, J. & Wei, L. The age-related changes in cartilage and osteoarthritis. Biomed Res Int 2013, 916530 (2013). 10.1155/2013/916530

54 Angelini, F. et al. Osteoarthritis endotype discovery via clustering of biochemical marker data. Ann Rheum Dis 81, 666–675 (2022). 10.1136/annrheumdis-2021-221763

55 Werdyani, S. et al. Endotypes of primary osteoarthritis identified by plasma metabolomics analysis. Rheumatology (Oxford) 60, 2735–2744 (2021). 10.1093/rheumatology/keaa693

56 Kühn, K. in Structure and Function of Collagen Types (eds Richard Mayne & Robert E. Burgeson) 1–42 (Academic Press, 1987).

57 Krane, S. M. The importance of proline residues in the structure, stability and susceptibility to proteolytic degradation of collagens. Amino Acids 35, 703–710 (2008). 10.1007/s00726-008-0073-2

58 Bailey, K. N. et al. Mechanosensitive Control of Articular Cartilage and Subchondral Bone Homeostasis in Mice Requires Osteocytic Transforming Growth Factor beta Signaling. Arthritis Rheumatol 73, 414–425 (2021). 10.1002/art.41548

59 Hu, Q. & Ecker, M. Overview of MMP-13 as a Promising Target for the Treatment of Osteoarthritis. Int J Mol Sci 22 (2021). 10.3390/ijms22041742

60 Bellis, S. L. Advantages of RGD peptides for directing cell association with biomaterials. Biomaterials 32, 4205–4210 (2011). 10.1016/j.biomaterials.2011.02.029

61 Sweeney, S. M. et al. Candidate cell and matrix interaction domains on the collagen fibril, the predominant protein of vertebrates. J Biol Chem 283, 21187–21197 (2008). 10.1074/jbc.M709319200

62 San Antonio, J. D., Jacenko, O., Fertala, A. & Orgel, J. Collagen Structure-Function Mapping Informs Applications for Regenerative Medicine. Bioengineering (Basel) 8 (2020). 10.3390/bioengineering8010003

63 Bachmann, M. F. & Oxenius, A. Interleukin 2: from immunostimulation to immunoregulation and back again. EMBO Rep 8, 1142–1148 (2007). 10.1038/sj.embor.7401099

64 Sondag, C. M. & Combs, C. K. Adhesion of monocytes to type I collagen stimulates an APP-dependent proinflammatory signaling response and release of Aβ1-40. J Neuroinflamm 7 (2010). 10.1186/1742-2094-7-22

65 George, A. & Hao, J. Role of phosphophoryn in dentin mineralization. Cells Tissues Organs 181, 232–240 (2005). 10.1159/000091384

66 Rappu, P., Salo, A. M., Myllyharju, J. & Heino, J. Role of prolyl hydroxylation in the molecular interactions of collagens. Essays Biochem 63, 325–335 (2019). 10.1042/EBC20180053

67 Garnero, P. The contribution of collagen crosslinks to bone strength. Bonekey Rep 1, 182 (2012). 10.1038/bonekey.2012.182

68 Grzesik, W. J. et al. Age-related changes in human bone proteoglycan structure. Impact of osteogenesis imperfecta. J Biol Chem 277, 43638–43647 (2002). 10.1074/jbc.M202124200

69 Chang, S. H. et al. Excessive mechanical loading promotes osteoarthritis through the gremlin-1-NF-kappaB pathway. Nat Commun 10, 1442 (2019). 10.1038/s41467-019-09491-5

70 Hosseininia, S. et al. Evidence for enhanced collagen type III deposition focally in the territorial matrix of osteoarthritic hip articular cartilage. Osteoarthritis Cartilage 24, 1029–1035 (2016). 10.1016/j.joca.2016.01.001

71 Volk, S. W. et al. Type III collagen regulates osteoblastogenesis and the quantity of trabecular bone. Calcif Tissue Int 94, 621–631 (2014). 10.1007/s00223-014-9843-x

72 Chiba, K., Ito, M., Osaki, M., Uetani, M. & Shindo, H. In vivo structural analysis of subchondral trabecular bone in osteoarthritis of the hip using multi-detector row CT. Osteoarthritis Cartilage 19, 180–185 (2011). 10.1016/j.joca.2010.11.002

73 Di Martino, A. et al. Collagen VI in the Musculoskeletal System. Int J Mol Sci 24 (2023). 10.3390/ijms24065095

74 Keene, D. R., Sakai, L. Y. & Burgeson, R. E. Human bone contains type III collagen, type VI collagen, and fibrillin: type III collagen is present on specific fibers that may mediate attachment of tendons, ligaments, and periosteum to calcified bone cortex. J Histochem Cytochem 39, 59–69 (1991). 10.1177/39.1.1983874

75 Williams, L., Layton, T., Yang, N., Feldmann, M. & Nanchahal, J. Collagen VI as a driver and disease biomarker in human fibrosis. FEBS J 289, 3603–3629 (2022). 10.1111/febs.16039

76 Komori, T. et al. Type VI Collagen Regulates Endochondral Ossification in the Temporomandibular Joint. JBMR Plus 6, e10617 (2022). 10.1002/jbm4.10617

77 Quarto, R., Dozin, B., Bonaldo, P., Cancedda, R. & Colombatti, A. Type VI collagen expression is upregulated in the early events of chondrocyte differentiation. Development 117, 245–251 (1993). 10.1242/dev.117.1.245

78 Zelenski, N. A. et al. Type VI Collagen Regulates Pericellular Matrix Properties, Chondrocyte Swelling, and Mechanotransduction in Mouse Articular Cartilage. Arthritis Rheumatol 67, 1286–1294 (2015). 10.1002/art.39034

79 Pullig, O., Weseloh, G. & Swoboda, B. Expression of type VI collagen in normal and osteoarthritic human cartilage. Osteoarthritis Cartilage 7, 191–202 (1999). 10.1053/joca.1998.0208

80 Williams SN, W. M., Bercovitz A. Hospitalization for total knee replacement among inpatients aged 45 and over: United States, 2000–2010. (2015). <https://www.cdc.gov/nchs/products/databriefs/db210.htm#:∼:text=For%20both%202000%20and%202010,women%20aged%2045%20and%20over.>.

81 Angel, P. M. et al. Mapping Extracellular Matrix Proteins in Formalin-Fixed, Paraffin-Embedded Tissues by MALDI Imaging Mass Spectrometry. J Proteome Res 17, 635–646 (2018). 10.1021/acs.jproteome.7b00713

82 Eckhard, U., Huesgen, P. F., Brandstetter, H. & Overall, C. M. Proteomic protease specificity profiling of clostridial collagenases reveals their intrinsic nature as dedicated degraders of collagen. J Proteomics 100, 102–114 (2014). 10.1016/j.jprot.2013.10.004

83 Eckhard, U., Schonauer, E., Nuss, D. & Brandstetter, H. Structure of collagenase G reveals a chew-and-digest mechanism of bacterial collagenolysis. Nat Struct Mol Biol 18, 1109–1114 (2011). 10.1038/nsmb.2127

84 Sauve, A. C. & Speed, T. P. Normalization, baseline correction and alignment of high-throughput mass spectrometry data. Proceedings Gensips, 1–4 (2004).

85 Deininger, S. O. et al. Normalization in MALDI-TOF imaging datasets of proteins: practical considerations. Anal Bioanal Chem 401, 167–181 (2011). 10.1007/s00216-011-4929-z

86 Trede, D. et al. Exploring three-dimensional matrix-assisted laser desorption/ionization imaging mass spectrometry data: three-dimensional spatial segmentation of mouse kidney. Anal Chem 84, 6079–6087 (2012). 10.1021/ac300673y

87 Zou, K. H., O’Malley, A. J. & Mauri, L. Receiver-operating characteristic analysis for evaluating diagnostic tests and predictive models. Circulation 115, 654–657 (2007). 10.1161/CIRCULATIONAHA.105.594929

88 Klein, O. et al. MALDI imaging mass spectrometry: discrimination of pathophysiological regions in traumatized skeletal muscle by characteristic peptide signatures. Proteomics 14, 2249–2260 (2014). 10.1002/pmic.201400088

89 Xia, J., Psychogios, N., Young, N. & Wishart, D. S. MetaboAnalyst: a web server for metabolomic data analysis and interpretation. Nucleic Acids Res 37, W652–660 (2009). 10.1093/nar/gkp356

90 Meier, F. et al. Online Parallel Accumulation-Serial Fragmentation (PASEF) with a Novel Trapped Ion Mobility Mass Spectrometer. Mol Cell Proteomics 17, 2534–2545 (2018). 10.1074/mcp.TIR118.000900

91 MacLean, B. X. et al. Using Skyline to Analyze Data-Containing Liquid Chromatography, Ion Mobility Spectrometry, and Mass Spectrometry Dimensions. J Am Soc Mass Spectrom 29, 2182–2188 (2018). 10.1007/s13361-018-2028-5

92 Tyanova, S., Temu, T. & Cox, J. The MaxQuant computational platform for mass spectrometry-based shotgun proteomics. Nat Protoc 11, 2301–2319 (2016). 10.1038/nprot.2016.136

