## Supplementary material for "Tissue and Extracellular Matrix Remodeling of the Subchondral Bone during Osteoarthritis of Knee Joints as revealed by Spatial Mass Spectrometry Imaging": Suppl Figure 1

### Slide 1
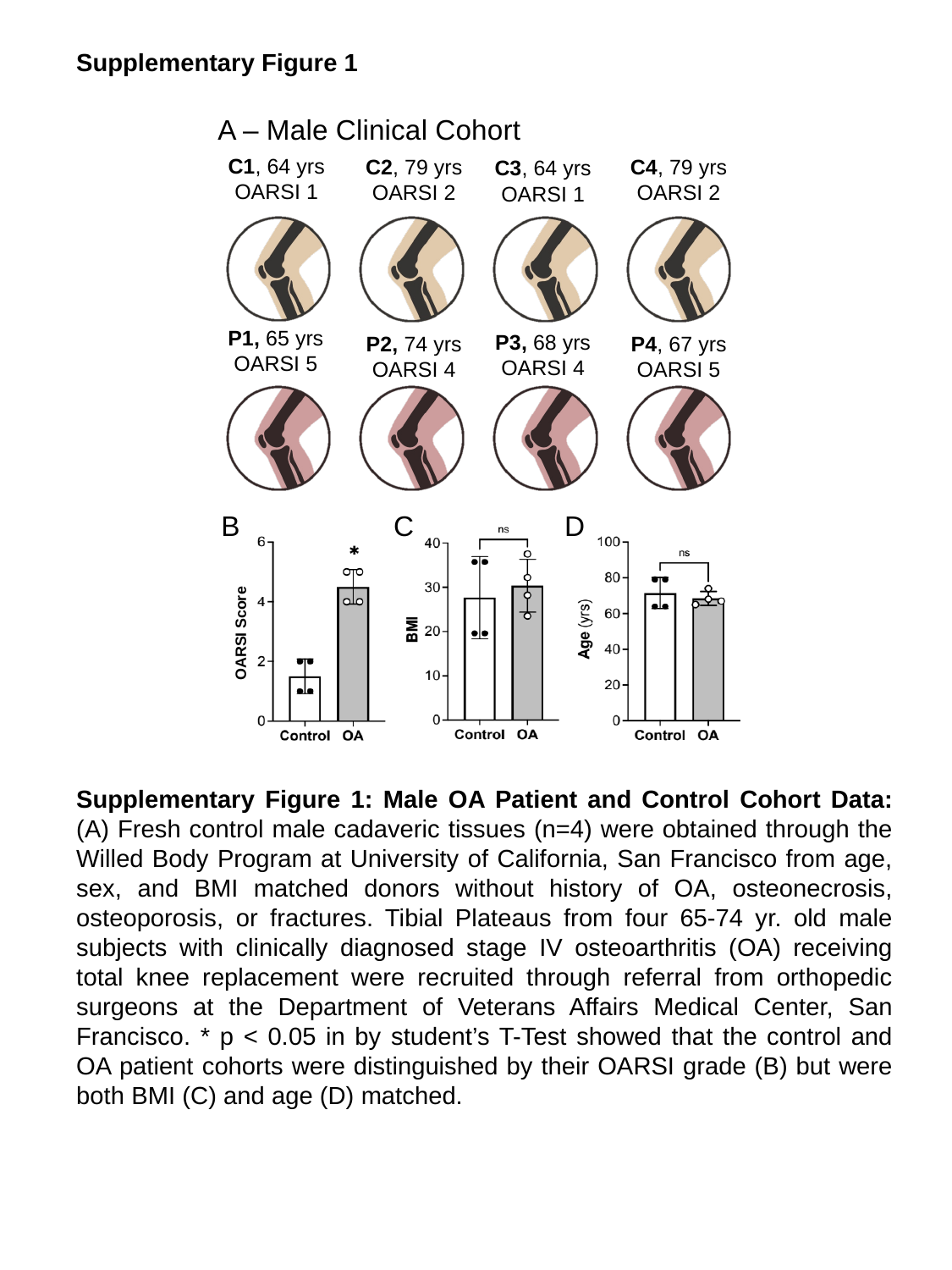

Supplementary Figure 1
A – Male Clinical Cohort
C1, 64 yrs
OARSI 1
C2, 79 yrs
OARSI 2
C4, 79 yrs
OARSI 2
C3, 64 yrs
OARSI 1
P1, 65 yrs
OARSI 5
P3, 68 yrs
OARSI 4
P2, 74 yrs
OARSI 4
P4, 67 yrs
OARSI 5
B
C
D
OARSI Score
Supplementary Figure 1: Male OA Patient and Control Cohort Data: (A) Fresh control male cadaveric tissues (n=4) were obtained through the Willed Body Program at University of California, San Francisco from age, sex, and BMI matched donors without history of OA, osteonecrosis, osteoporosis, or fractures. Tibial Plateaus from four 65-74 yr. old male subjects with clinically diagnosed stage IV osteoarthritis (OA) receiving total knee replacement were recruited through referral from orthopedic surgeons at the Department of Veterans Affairs Medical Center, San Francisco. * p < 0.05 in by student’s T-Test showed that the control and OA patient cohorts were distinguished by their OARSI grade (B) but were both BMI (C) and age (D) matched.
