## Supplementary material for "Tissue and Extracellular Matrix Remodeling of the Subchondral Bone during Osteoarthritis of Knee Joints as revealed by Spatial Mass Spectrometry Imaging": Suppl Figure 2

### Supplemental Figure 2

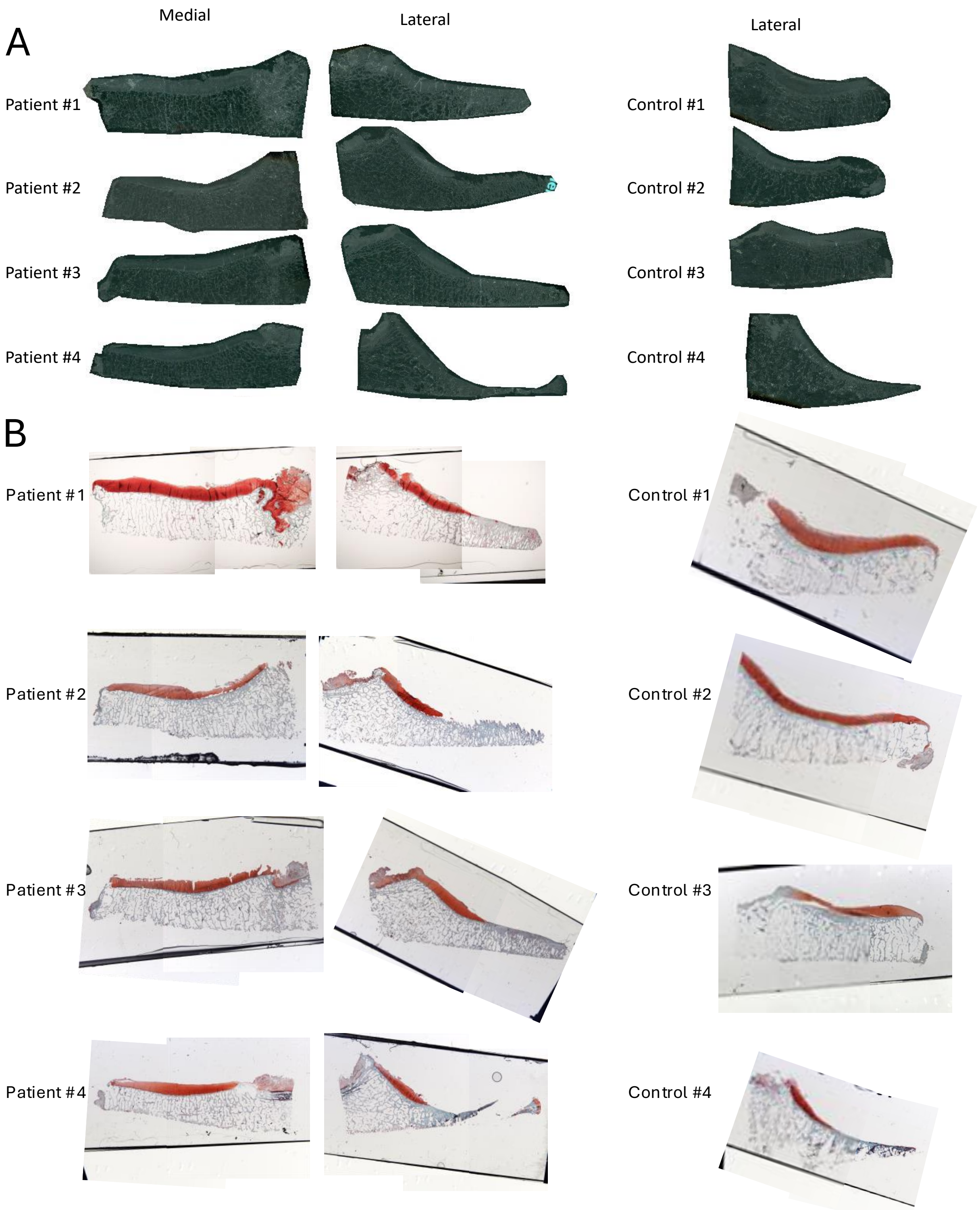


**OA & Control Tissue Sections and Safranin O Staining.** (A) Unstained tissue scans and (B) Safranin-O Histologic staining of FFPE slides of medial and lateral halves of the Tibial Plateau of TKA Patients and the lateral half from donor cadaveric non-arthritic Controls. Safranin O staining appeared as red to stain proteoglycans in cartilage layers while the subchondral bone remained white/grey.
