## Supplementary material for "Tissue and Extracellular Matrix Remodeling of the Subchondral Bone during Osteoarthritis of Knee Joints as revealed by Spatial Mass Spectrometry Imaging": Suppl Figure 3

### Slide 1
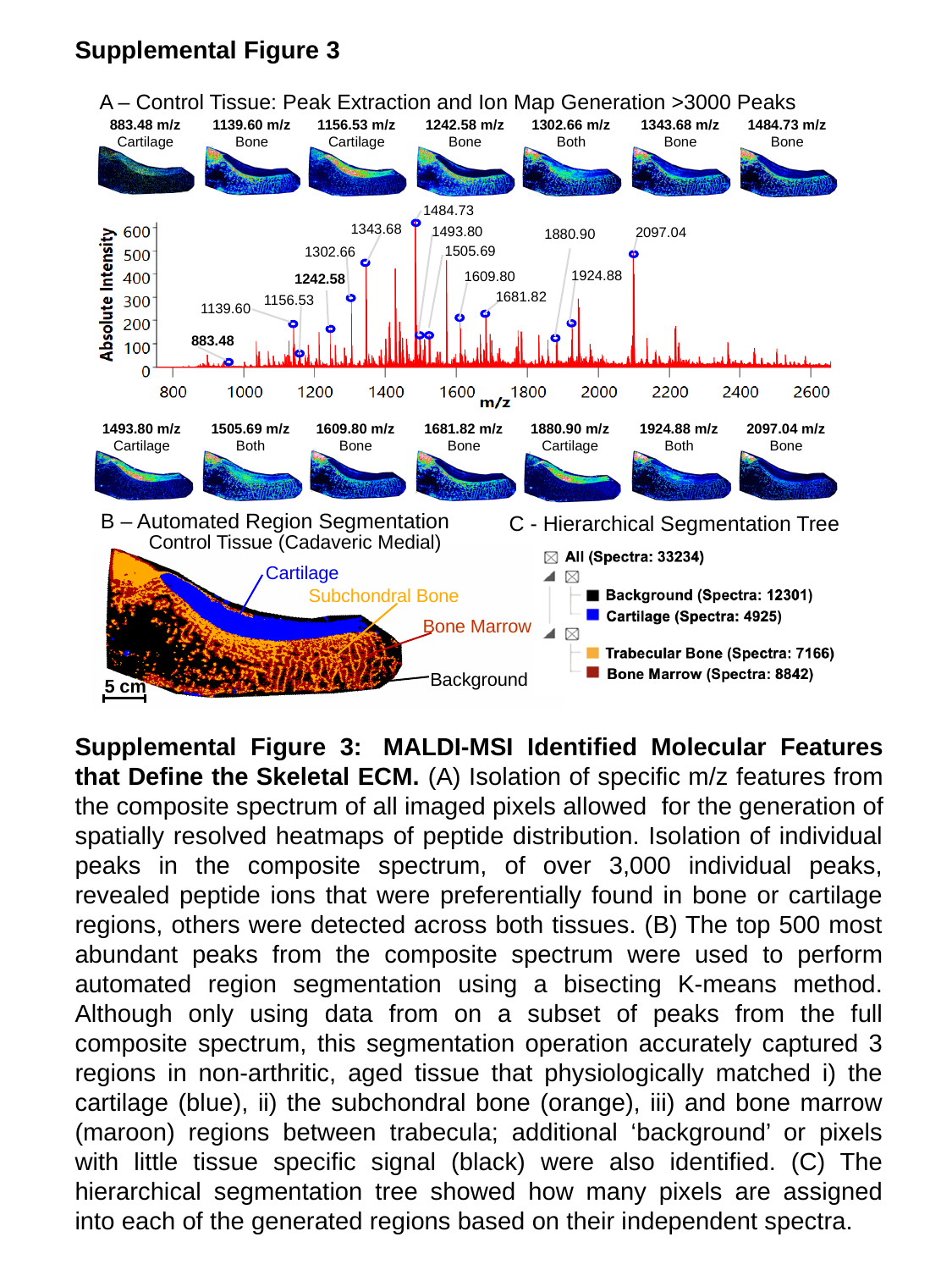

Supplemental Figure 3
A – Control Tissue: Peak Extraction and Ion Map Generation >3000 Peaks
883.48 m/z
Cartilage
1139.60 m/z
Bone
1156.53 m/z
Cartilage
1242.58 m/z
Bone
1302.66 m/z
Both
1343.68 m/z
Bone
1484.73 m/z
Bone
1484.73
1343.68
1493.80
2097.04
1880.90
1505.69
1302.66
1924.88
1609.80
1242.58
1681.82
1156.53
1139.60
883.48
1493.80 m/z
Cartilage
1505.69 m/z
Both
1609.80 m/z
Bone
1681.82 m/z
Bone
1880.90 m/z
Cartilage
1924.88 m/z
Both
2097.04 m/z
Bone
B – Automated Region Segmentation
C - Hierarchical Segmentation Tree
Control Tissue (Cadaveric Medial)
Cartilage
Subchondral Bone
Bone Marrow
Background
5 cm
Supplemental Figure 3:  MALDI-MSI Identified Molecular Features that Define the Skeletal ECM. (A) Isolation of specific m/z features from the composite spectrum of all imaged pixels allowed  for the generation of spatially resolved heatmaps of peptide distribution. Isolation of individual peaks in the composite spectrum, of over 3,000 individual peaks, revealed peptide ions that were preferentially found in bone or cartilage regions, others were detected across both tissues. (B) The top 500 most abundant peaks from the composite spectrum were used to perform automated region segmentation using a bisecting K-means method. Although only using data from on a subset of peaks from the full composite spectrum, this segmentation operation accurately captured 3 regions in non-arthritic, aged tissue that physiologically matched i) the cartilage (blue), ii) the subchondral bone (orange), iii) and bone marrow (maroon) regions between trabecula; additional ‘background’ or pixels with little tissue specific signal (black) were also identified. (C) The hierarchical segmentation tree showed how many pixels are assigned into each of the generated regions based on their independent spectra.
