## Supplementary material for "Tissue and Extracellular Matrix Remodeling of the Subchondral Bone during Osteoarthritis of Knee Joints as revealed by Spatial Mass Spectrometry Imaging": Suppl Figure 4

### Slide 1
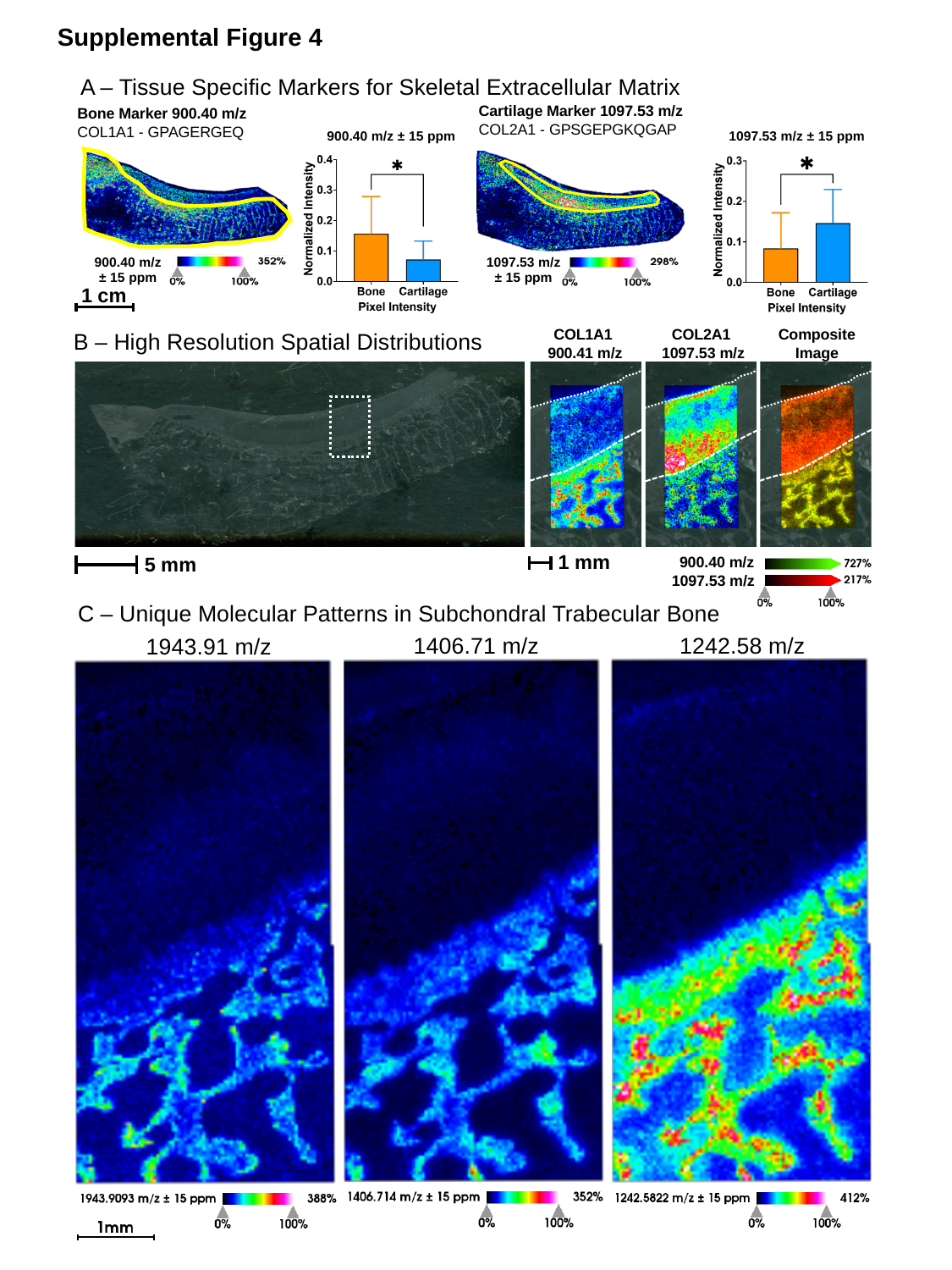

Supplemental Figure 4
A – Tissue Specific Markers for Skeletal Extracellular Matrix
Cartilage Marker 1097.53 m/z
COL2A1 - GPSGEPGKQGAP
Bone Marker 900.40 m/z
COL1A1 - GPAGERGEQ
900.40 m/z ± 15 ppm
1097.53 m/z ± 15 ppm
900.40 m/z
± 15 ppm
1097.53 m/z
± 15 ppm
1 cm
COL1A1
900.41 m/z
COL2A1
1097.53 m/z
Composite
Image
B – High Resolution Spatial Distributions
1 mm
 900.40 m/z
5 mm
1097.53 m/z
C – Unique Molecular Patterns in Subchondral Trabecular Bone
1406.71 m/z
1242.58 m/z
1943.91 m/z

### Slide 2
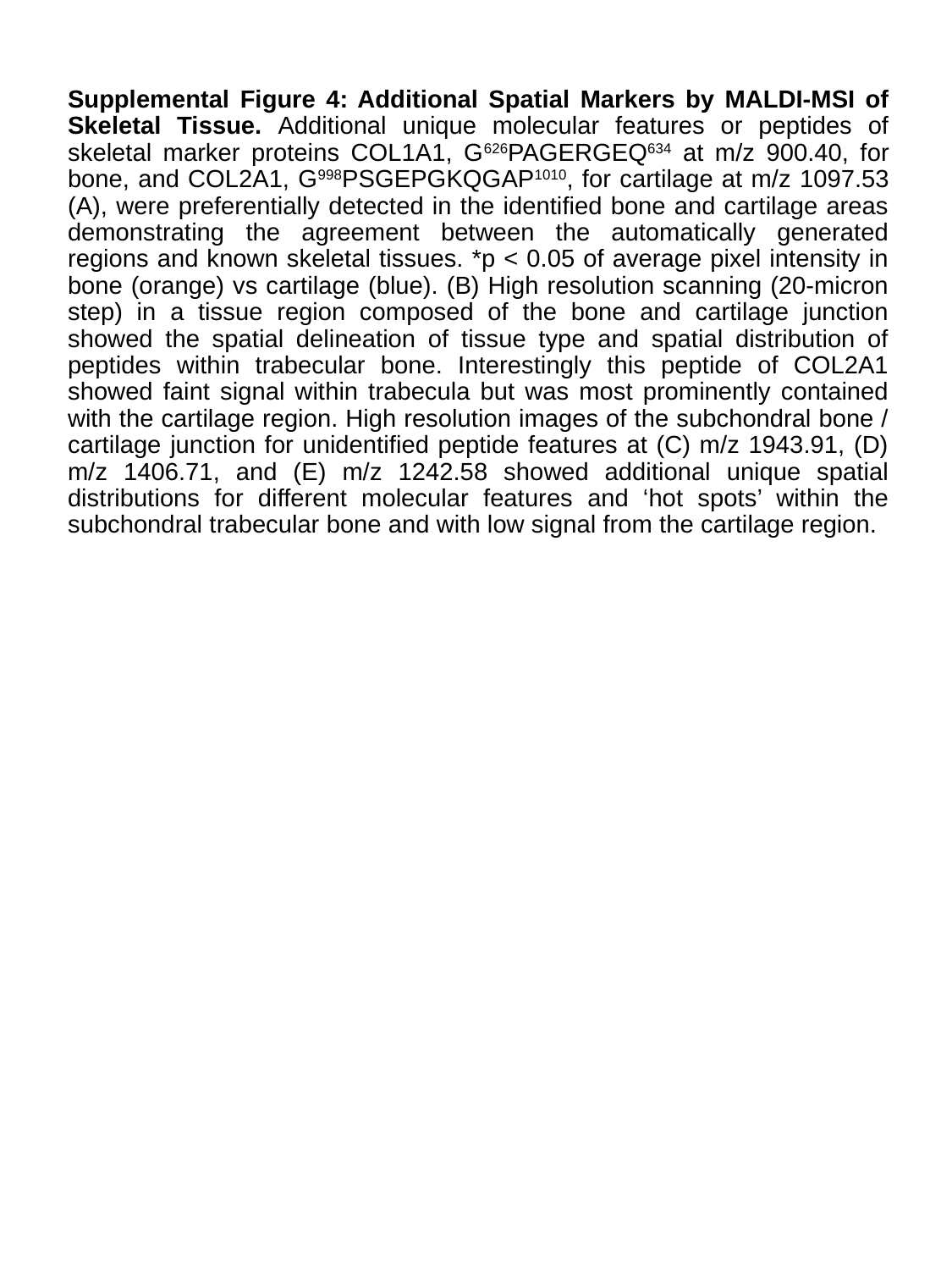

Supplemental Figure 4: Additional Spatial Markers by MALDI-MSI of Skeletal Tissue. Additional unique molecular features or peptides of skeletal marker proteins COL1A1, G^626^PAGERGEQ^634^ at m/z 900.40, for bone, and COL2A1, G^998^PSGEPGKQGAP^1010^, for cartilage at m/z 1097.53 (A), were preferentially detected in the identified bone and cartilage areas demonstrating the agreement between the automatically generated regions and known skeletal tissues. *p < 0.05 of average pixel intensity in bone (orange) vs cartilage (blue). (B) High resolution scanning (20-micron step) in a tissue region composed of the bone and cartilage junction showed the spatial delineation of tissue type and spatial distribution of peptides within trabecular bone. Interestingly this peptide of COL2A1 showed faint signal within trabecula but was most prominently contained with the cartilage region. High resolution images of the subchondral bone / cartilage junction for unidentified peptide features at (C) m/z 1943.91, (D) m/z 1406.71, and (E) m/z 1242.58 showed additional unique spatial distributions for different molecular features and ‘hot spots’ within the subchondral trabecular bone and with low signal from the cartilage region.
