## Supplementary material for "Tissue and Extracellular Matrix Remodeling of the Subchondral Bone during Osteoarthritis of Knee Joints as revealed by Spatial Mass Spectrometry Imaging": Suppl Figure 5

### Slide 1
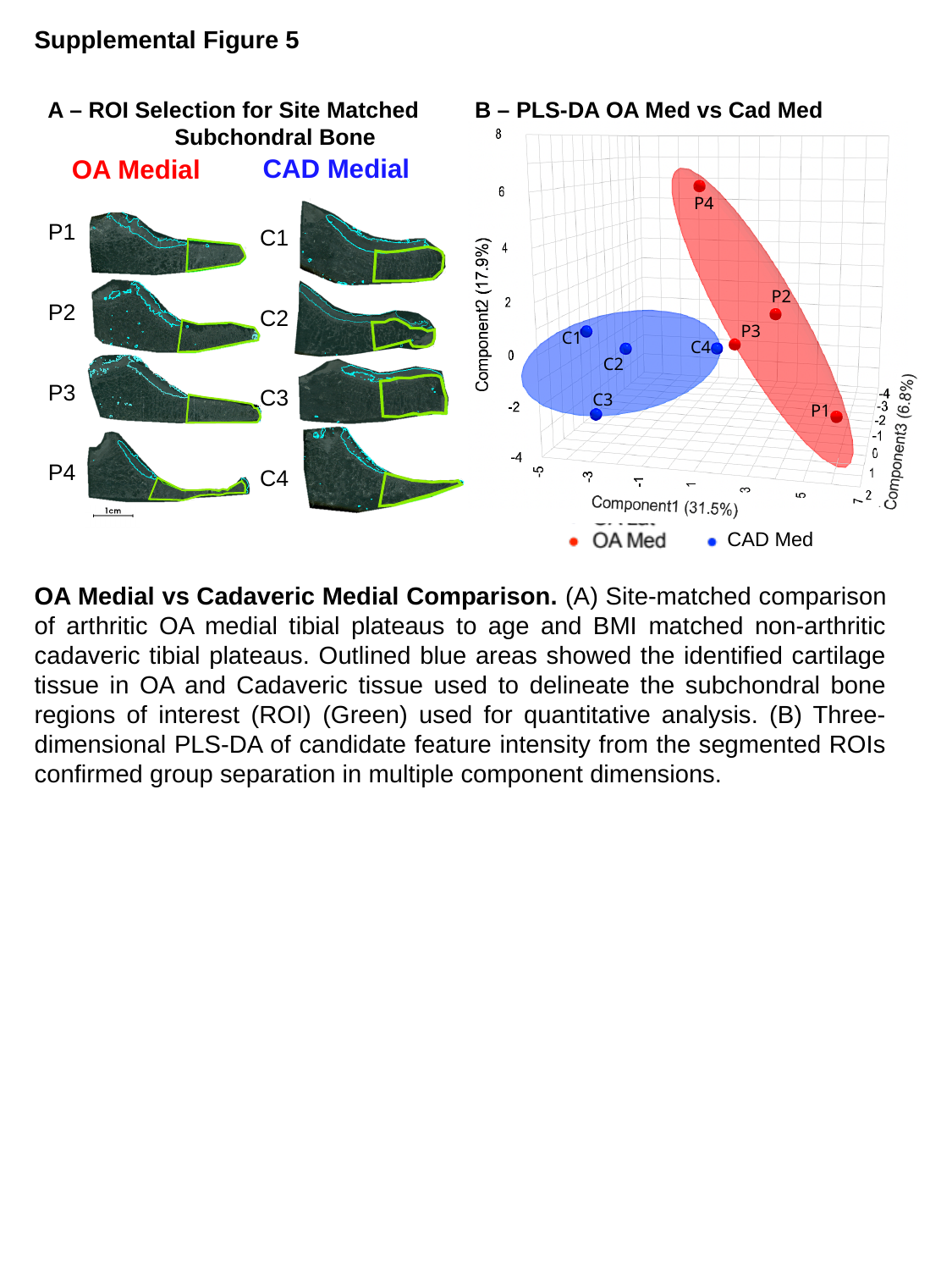

Supplemental Figure 5
B – PLS-DA OA Med vs Cad Med
A – ROI Selection for Site Matched 	Subchondral Bone
P4
P2
P3
C1
C4
C2
C3
P1
CAD Medial
OA Medial
C1
C2
C3
C4
P1
P2
P3
P4
CAD Med
OA Medial vs Cadaveric Medial Comparison. (A) Site-matched comparison of arthritic OA medial tibial plateaus to age and BMI matched non-arthritic cadaveric tibial plateaus. Outlined blue areas showed the identified cartilage tissue in OA and Cadaveric tissue used to delineate the subchondral bone regions of interest (ROI) (Green) used for quantitative analysis. (B) Three-dimensional PLS-DA of candidate feature intensity from the segmented ROIs confirmed group separation in multiple component dimensions.
