## Supplementary material for "Tissue and Extracellular Matrix Remodeling of the Subchondral Bone during Osteoarthritis of Knee Joints as revealed by Spatial Mass Spectrometry Imaging": Suppl Figure 6

### Supplemental Figure 6

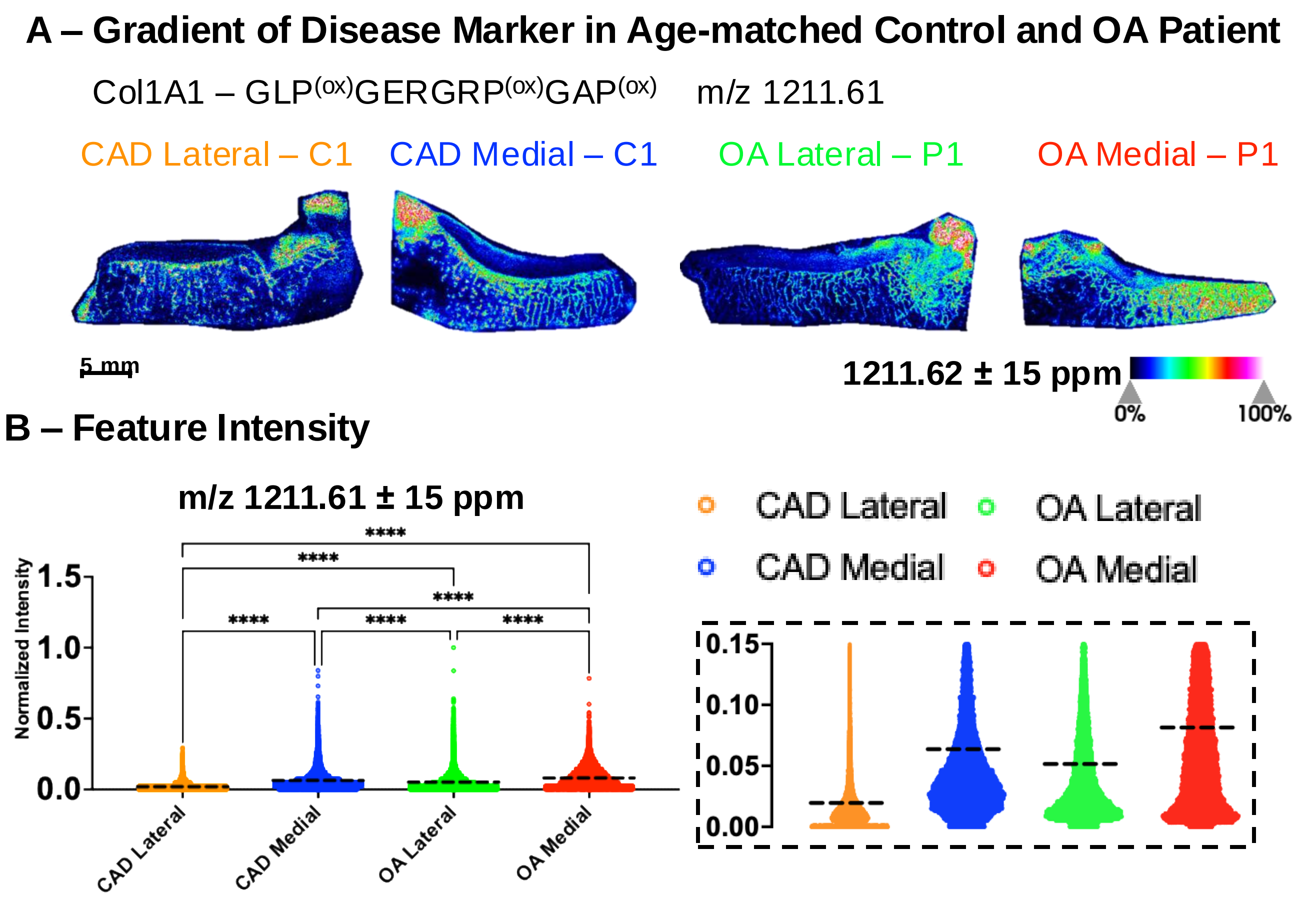


**Gradient of Disease in Age-related Osteoarthritis.** A) Heatmap for a candidate peptide, GLP$^{(ox)}$GERGRP$^{(ox)}$GAP$^{(ox)}$ at m/z 1211.61, from Collagen alpha-1(I) (Col1A1) that was found upregulated in OA is depicted across both halves of Cadaveric control C1 and OA patient P1 tissue sections. B) Quantification of normalized pixel intensity for the same peptide showed a gradient of increase from Cadaveric Lateral to OA Medial tissue sections. All four samples were statistically different from one another by post-hoc Tukey testing after one-way ANOVA ($p<0.05$). Inset (dotted box) showed increased resolution on the y-axis (between 0 and 0.15) around the mean pixel intensity values for each tissue section to more precisely depict differences in feature abundance across the four tissue sections.
