## Supplementary material for "Tissue and Extracellular Matrix Remodeling of the Subchondral Bone during Osteoarthritis of Knee Joints as revealed by Spatial Mass Spectrometry Imaging": Suppl Figure 7

### Slide 1
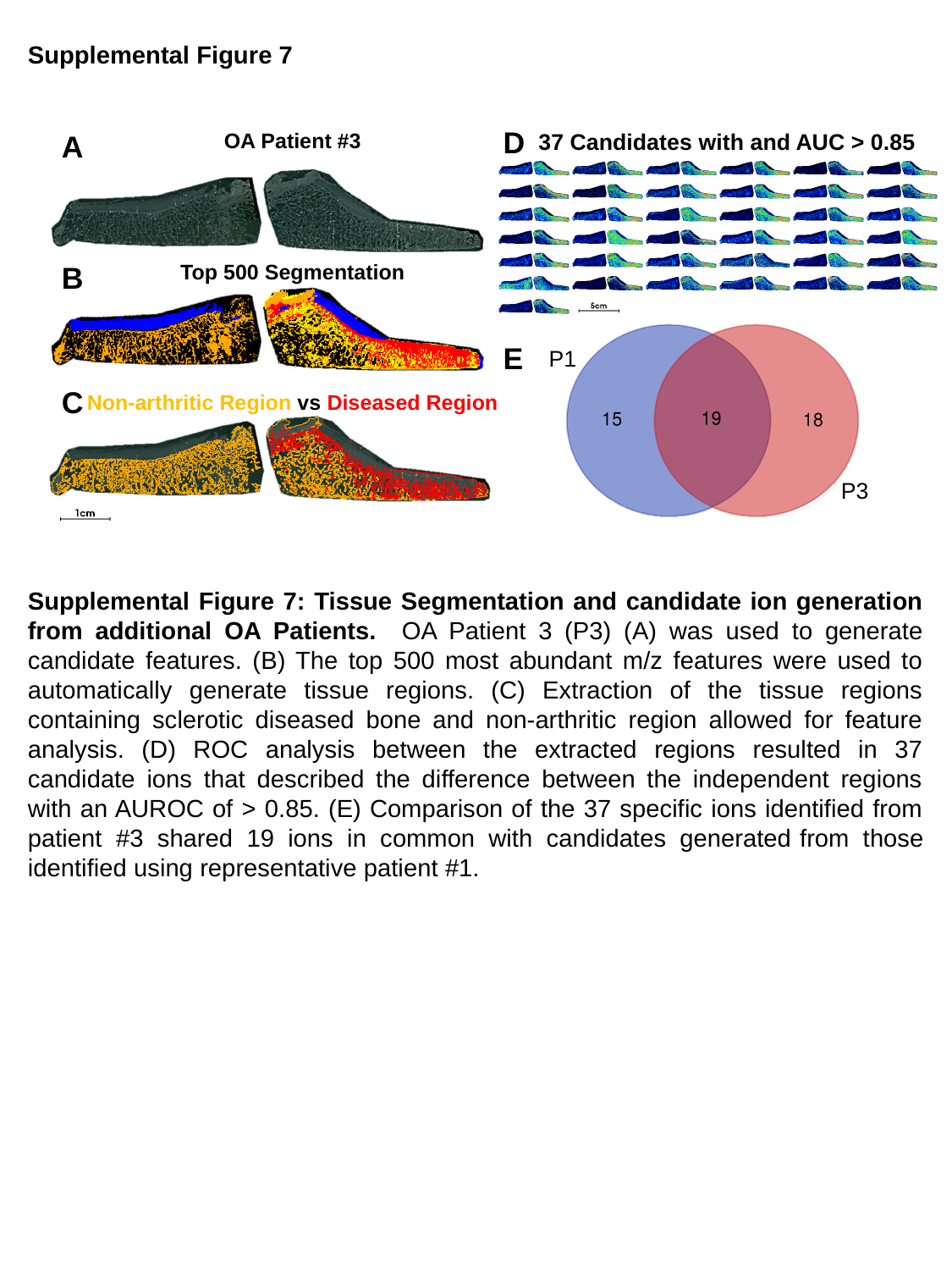

Supplemental Figure 7
D
A
OA Patient #3
37 Candidates with and AUC > 0.85
B
Top 500 Segmentation
E
P1
C
Non-arthritic Region vs Diseased Region
P3
Supplemental Figure 7: Tissue Segmentation and candidate ion generation from additional OA Patients. OA Patient 3 (P3) (A) was used to generate candidate features. (B) The top 500 most abundant m/z features were used to automatically generate tissue regions. (C) Extraction of the tissue regions containing sclerotic diseased bone and non-arthritic region allowed for feature analysis. (D) ROC analysis between the extracted regions resulted in 37 candidate ions that described the difference between the independent regions with an AUROC of > 0.85. (E) Comparison of the 37 specific ions identified from patient #3 shared 19 ions in common with candidates generated from those identified using representative patient #1.
