## Supplementary material for "Tissue and Extracellular Matrix Remodeling of the Subchondral Bone during Osteoarthritis of Knee Joints as revealed by Spatial Mass Spectrometry Imaging": Suppl Figure 8

### Slide 1
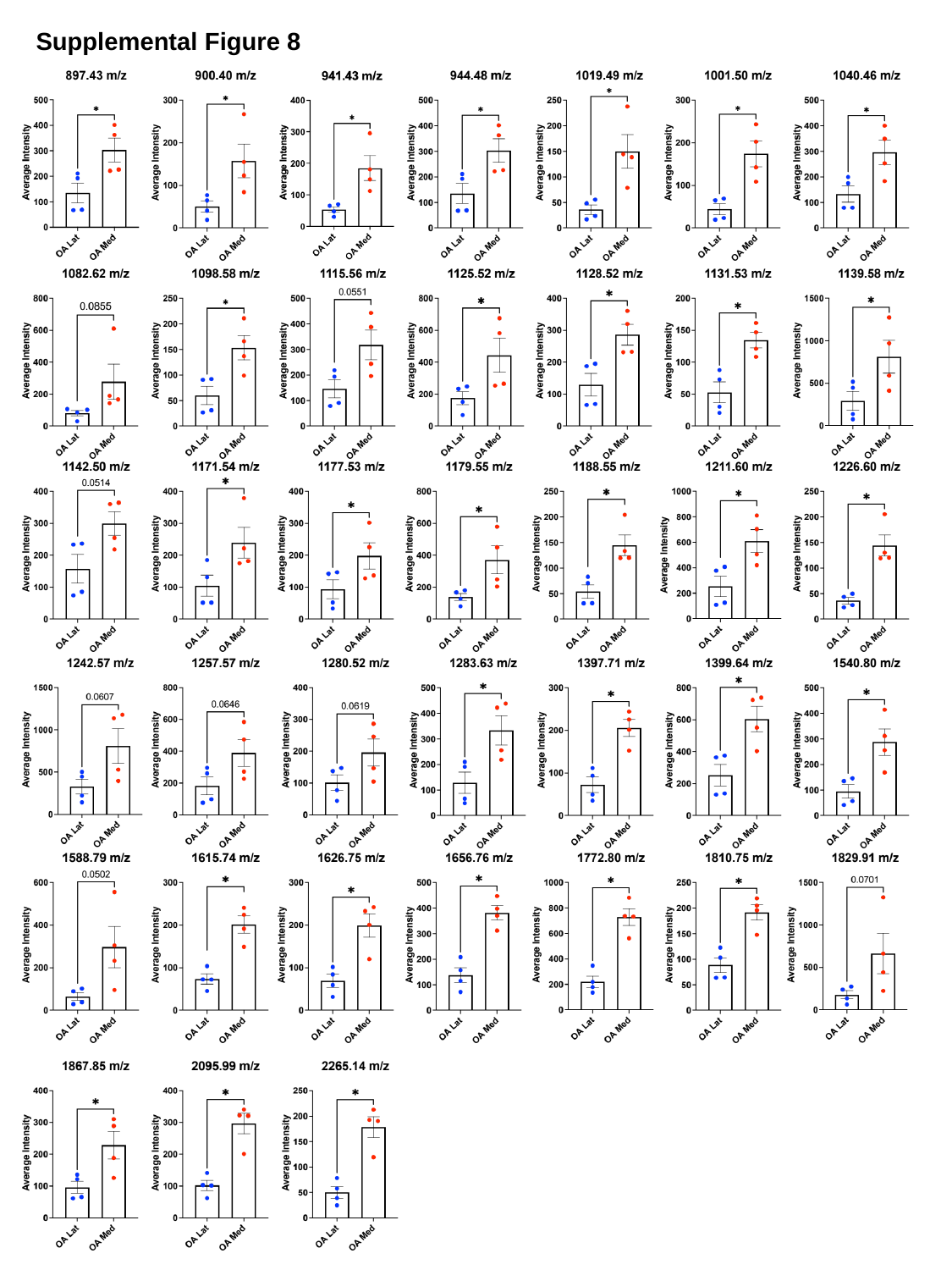

Supplemental Figure 8

### Slide 2
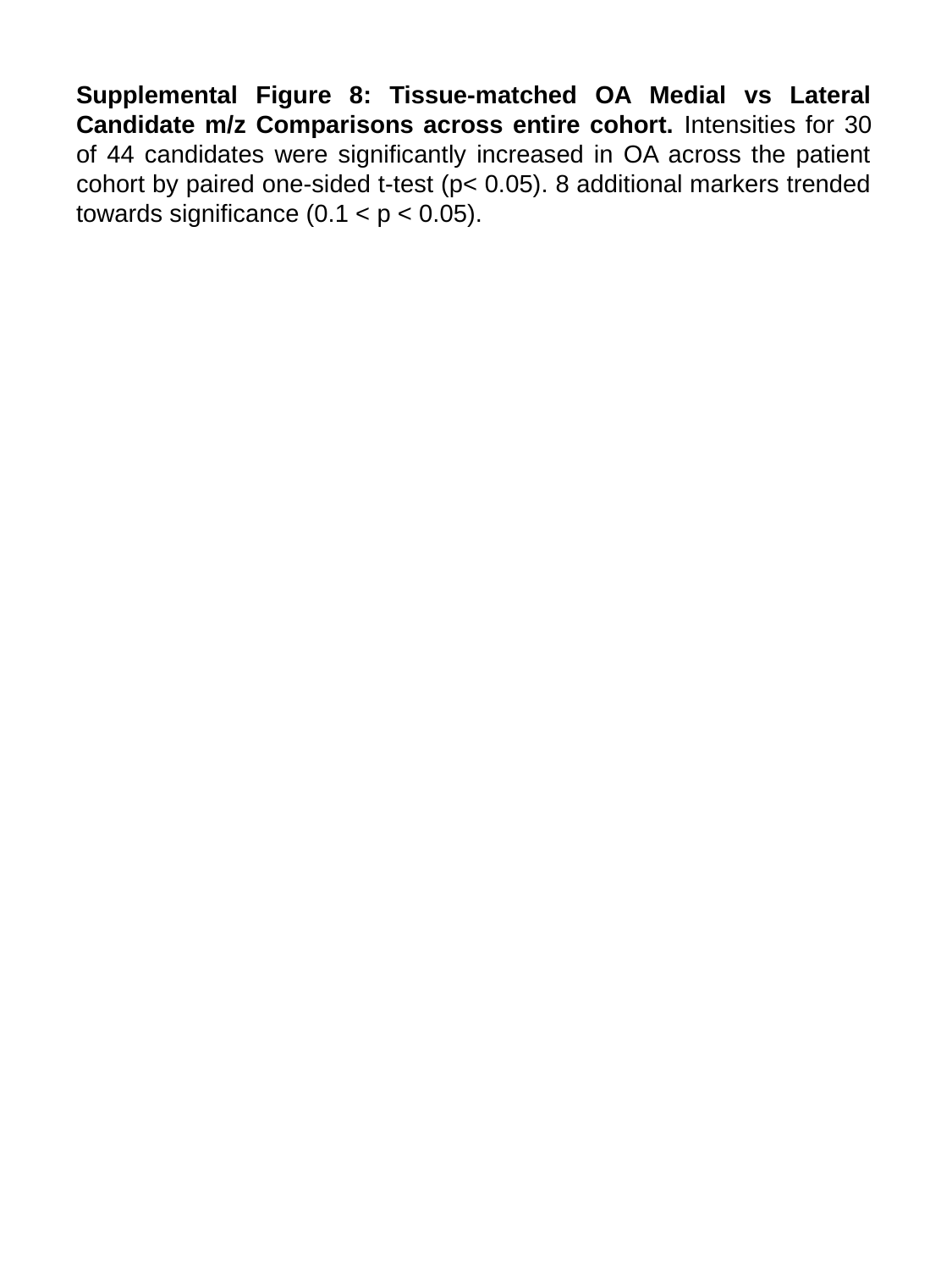

#### Supplemental Figure 8: Tissue-matched OA Medial vs Lateral Candidate m/z Comparisons across entire cohort. Intensities for 30 of 44 candidates were significantly increased in OA across the patient cohort by paired one-sided t-test (p< 0.05). 8 additional markers trended towards significance (0.1 < p < 0.05).
