## Supplementary material for "Tissue and Extracellular Matrix Remodeling of the Subchondral Bone during Osteoarthritis of Knee Joints as revealed by Spatial Mass Spectrometry Imaging": Suppl Figure 9

### Slide 1
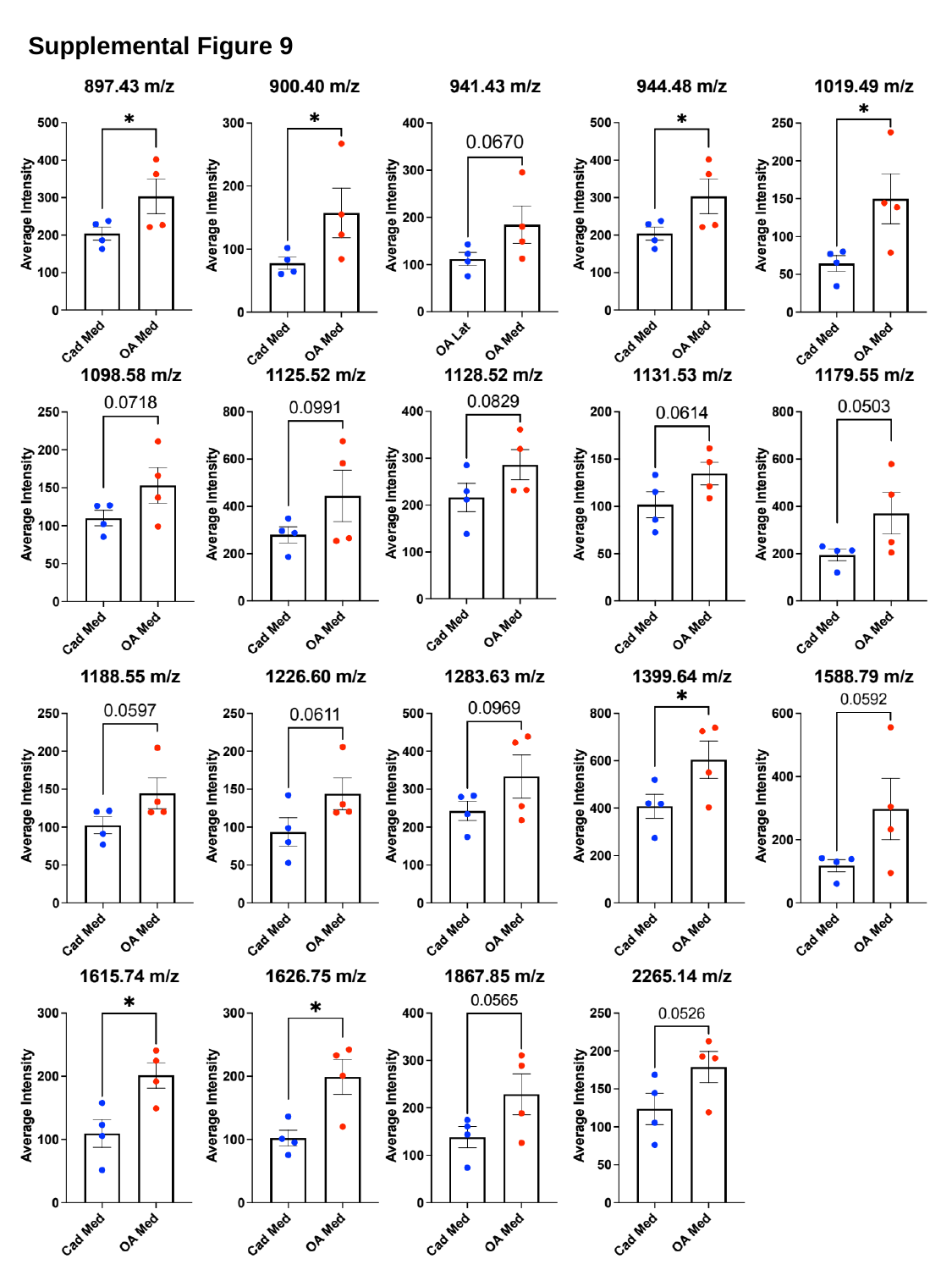

Supplemental Figure 9

### Slide 2
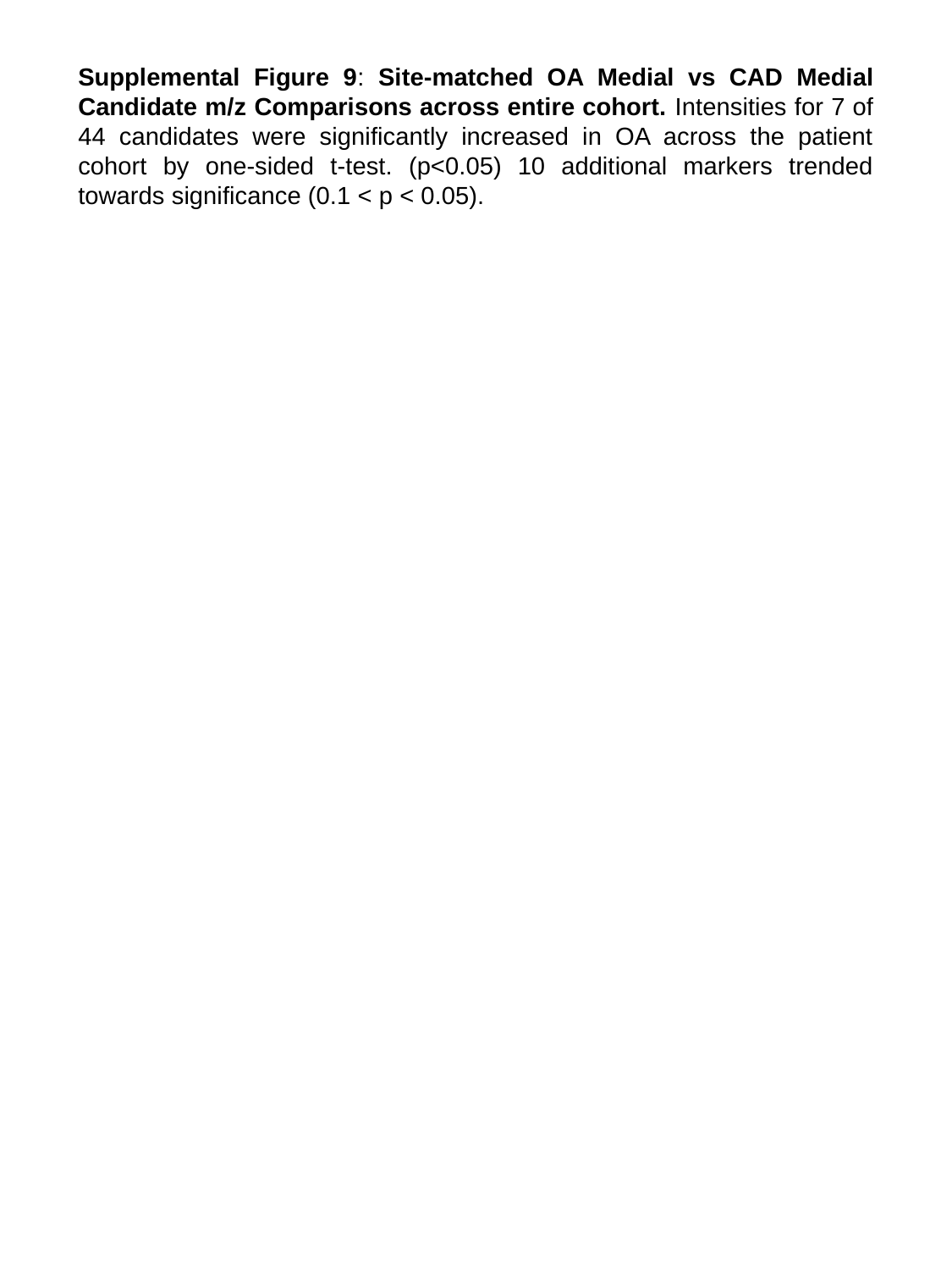

Supplemental Figure 9: Site-matched OA Medial vs CAD Medial Candidate m/z Comparisons across entire cohort. Intensities for 7 of 44 candidates were significantly increased in OA across the patient cohort by one-sided t-test. (p<0.05) 10 additional markers trended towards significance (0.1 < p < 0.05).
