## Supplementary material for "Tissue and Extracellular Matrix Remodeling of the Subchondral Bone during Osteoarthritis of Knee Joints as revealed by Spatial Mass Spectrometry Imaging": Suppl Figure 10

### Slide 1
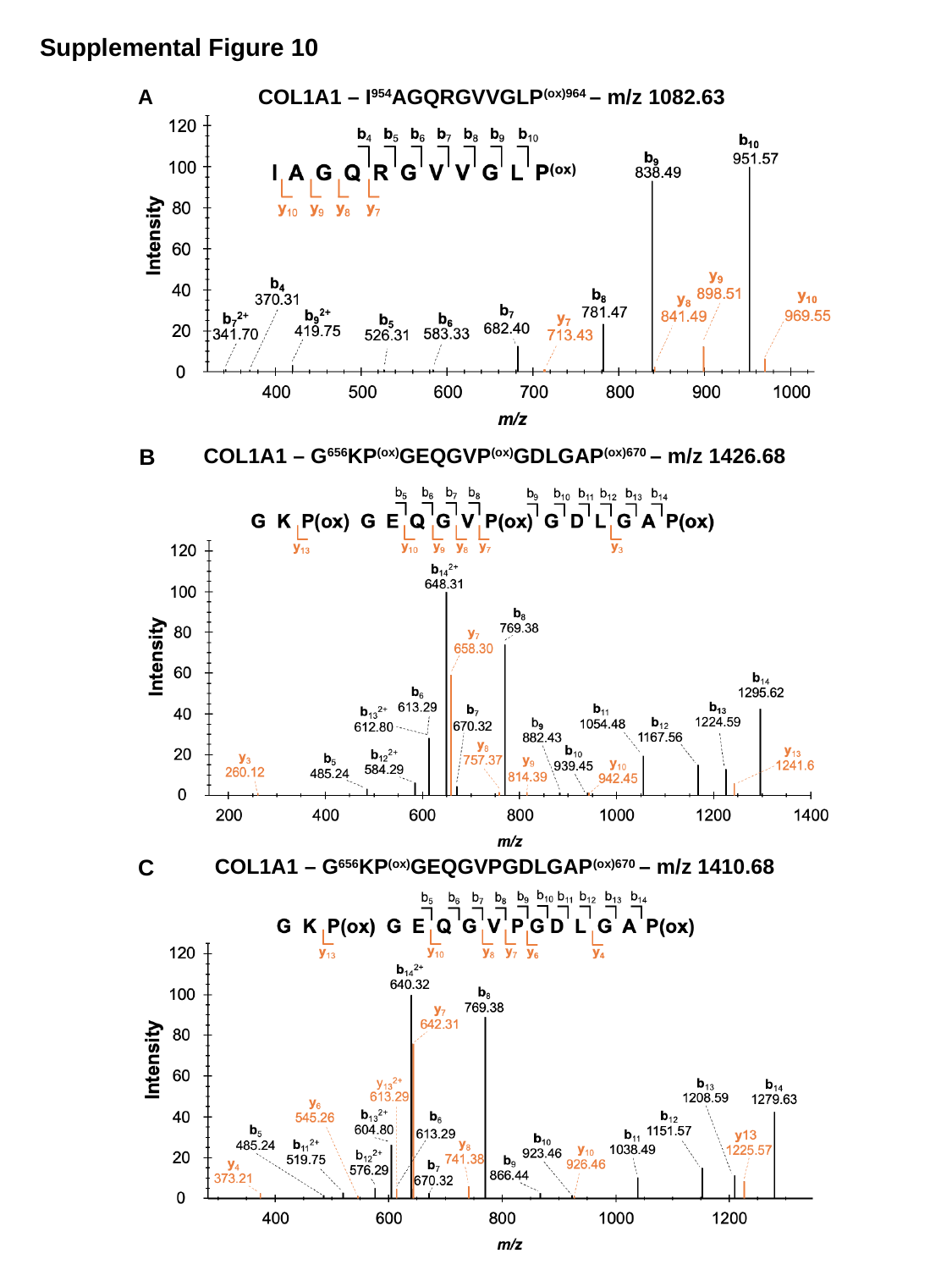

Supplemental Figure 10
A
COL1A1 – I^954^AGQRGVVGLP^(ox)964^ – m/z 1082.63
B
COL1A1 – G^656^KP^(ox)^GEQGVP^(ox)^GDLGAP^(ox)670^ – m/z 1426.68
C
COL1A1 – G^656^KP^(ox)^GEQGVPGDLGAP^(ox)670^ – m/z 1410.68

### Slide 2
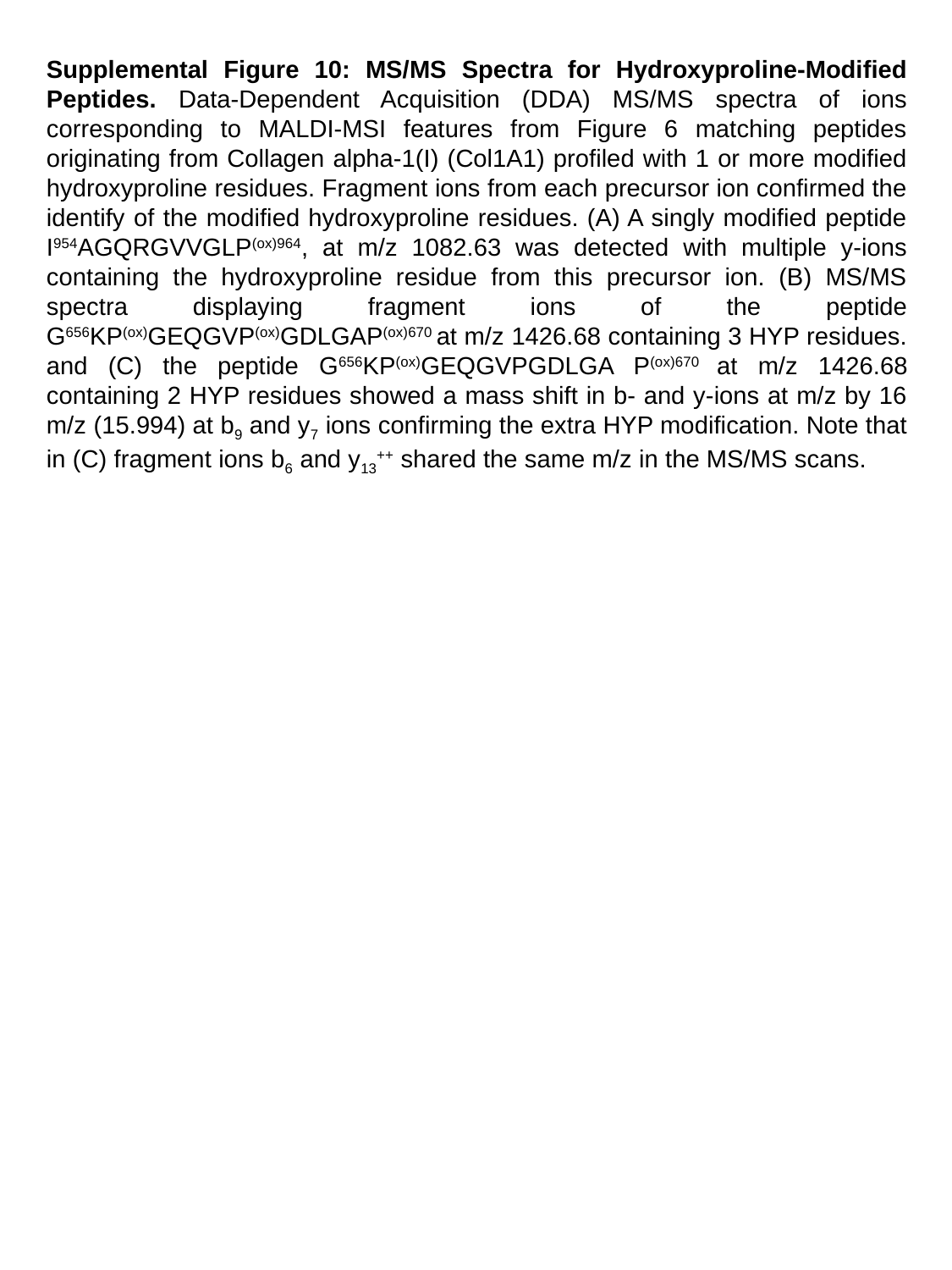

Supplemental Figure 10: MS/MS Spectra for Hydroxyproline-Modified Peptides. Data-Dependent Acquisition (DDA) MS/MS spectra of ions corresponding to MALDI-MSI features from Figure 6 matching peptides originating from Collagen alpha-1(I) (Col1A1) profiled with 1 or more modified hydroxyproline residues. Fragment ions from each precursor ion confirmed the identify of the modified hydroxyproline residues. (A) A singly modified peptide I^954^AGQRGVVGLP^(ox)964^, at m/z 1082.63 was detected with multiple y-ions containing the hydroxyproline residue from this precursor ion. (B) MS/MS spectra displaying fragment ions of the peptide G^656^KP^(ox)^GEQGVP^(ox)^GDLGAP^(ox)670^ at m/z 1426.68 containing 3 HYP residues. and (C) the peptide G^656^KP^(ox)^GEQGVPGDLGA P^(ox)670^ at m/z 1426.68 containing 2 HYP residues showed a mass shift in b- and y-ions at m/z by 16 m/z (15.994) at b_9_ and y_7_ ions confirming the extra HYP modification. Note that in (C) fragment ions b_6_ and y_13_^++^ shared the same m/z in the MS/MS scans.
