## Supplementary material for "Tissue and Extracellular Matrix Remodeling of the Subchondral Bone during Osteoarthritis of Knee Joints as revealed by Spatial Mass Spectrometry Imaging": Suppl Figure 11

### Slide 1
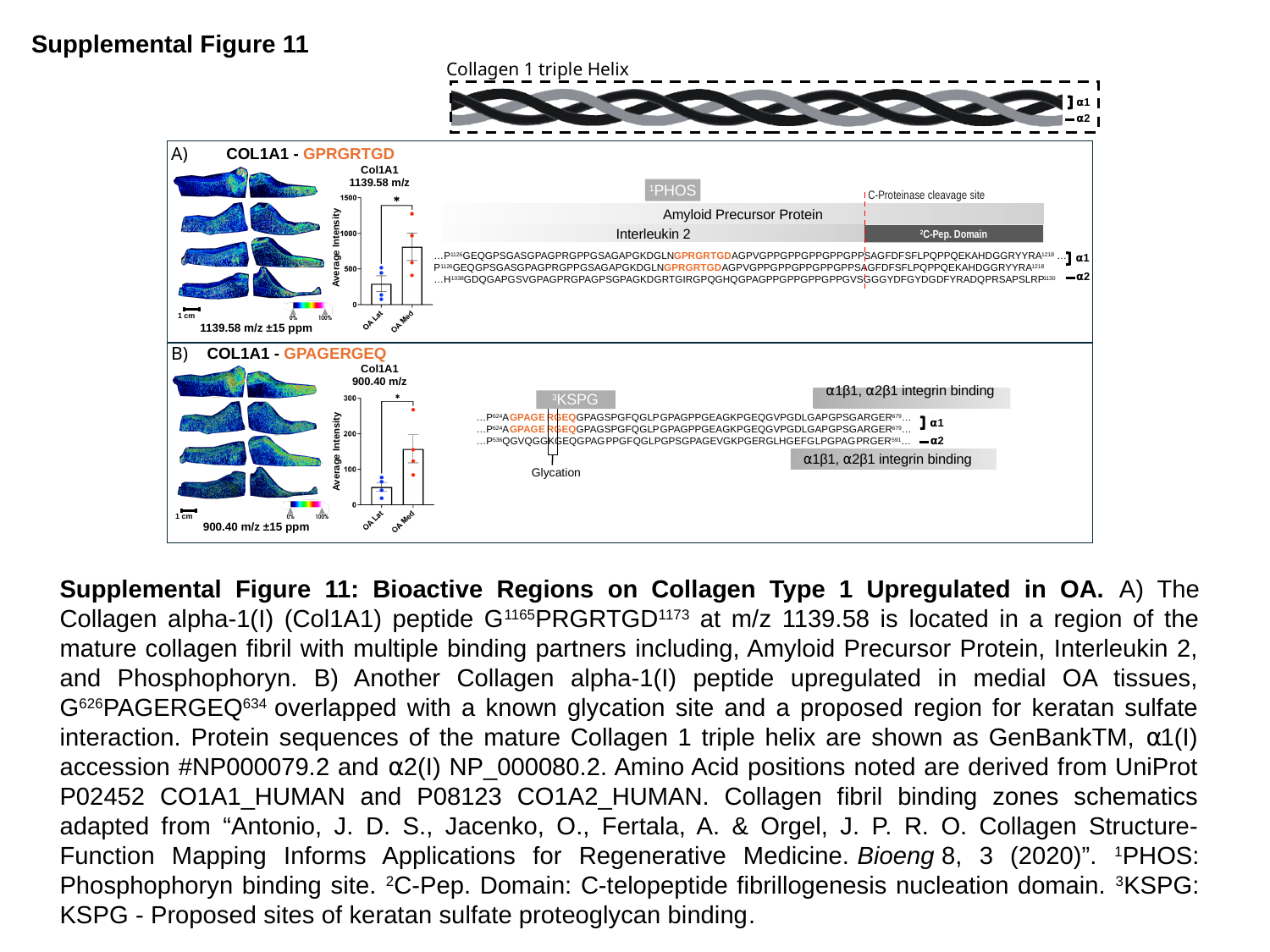

Supplemental Figure 11
Collagen 1 triple Helix
⍺1
⍺2
A)
COL1A1 - GPRGRTGD
Col1A1 1139.58 m/z
^1^PHOS
C-Proteinase cleavage site
Average Intensity
Amyloid Precursor Protein
Interleukin 2
^2^C-Pep. Domain
…P^1126^GEQGPSGASGPAGPRGPPGSAGAPGKDGLNGPRGRTGDAGPVGPPGPPGPPGPPGPPSAGFDFSFLPQPPQEKAHDGGRYYRA^1218^ …P^1126^GEQGPSGASGPAGPRGPPGSAGAPGKDGLNGPRGRTGDAGPVGPPGPPGPPGPPGPPSAGFDFSFLPQPPQEKAHDGGRYYRA^1218^
…H^1038^GDQGAPGSVGPAGPRGPAGPSGPAGKDGRTGIRGPQGHQGPAGPPGPPGPPGPPGVSGGGYDFGYDGDFYRADQPRSAPSLRP^1130^
⍺1
⍺2
1 cm
1139.58 m/z ±15 ppm
B)
COL1A1 - GPAGERGEQ
Col1A1 900.40 m/z
^3^KSPG
Average Intensity
⍺1β1, ⍺2β1 integrin binding
…P^624^A GPAGE RGEQGPAGSPGFQGLP GPAGPPGEAGKPGEQGVPGDLGAPGPSGARGER^679^…
…P^624^A GPAGE RGEQGPAGSPGFQGLP GPAGPPGEAGKPGEQGVPGDLGAPGPSGARGER^679^…
…P^536^QGVQGGKGEQGPAG PPGFQGLPGPSGPAGEVGKPGERGLHGEFG LPGPAG PRGER^591^…
⍺1
⍺2
⍺1β1, ⍺2β1 integrin binding
Glycation
1 cm
900.40 m/z ±15 ppm
Supplemental Figure 11: Bioactive Regions on Collagen Type 1 Upregulated in OA. A) The Collagen alpha-1(I) (Col1A1) peptide G^1165^PRGRTGD^1173^ at m/z 1139.58 is located in a region of the mature collagen fibril with multiple binding partners including, Amyloid Precursor Protein, Interleukin 2, and Phosphophoryn. B) Another Collagen alpha-1(I) peptide upregulated in medial OA tissues, G^626^PAGERGEQ^634^ overlapped with a known glycation site and a proposed region for keratan sulfate interaction. Protein sequences of the mature Collagen 1 triple helix are shown as GenBankTM, ⍺1(I) accession #NP000079.2 and ⍺2(I) NP_000080.2. Amino Acid positions noted are derived from UniProt P02452 CO1A1_HUMAN and P08123 CO1A2_HUMAN. Collagen fibril binding zones schematics adapted from “Antonio, J. D. S., Jacenko, O., Fertala, A. & Orgel, J. P. R. O. Collagen Structure-Function Mapping Informs Applications for Regenerative Medicine. Bioeng 8, 3 (2020)”. ^1^PHOS: Phosphophoryn binding site. ^2^C-Pep. Domain: C-telopeptide fibrillogenesis nucleation domain. ^3^KSPG: KSPG - Proposed sites of keratan sulfate proteoglycan binding.
